# Reduced purine biosynthesis in humans after their divergence from Neandertals

**DOI:** 10.1101/2020.05.11.087338

**Authors:** Vita Stepanova, Kaja Ewa Moczulska, Guido N. Vacano, Xiang-chun Ju, Stephan Riesenberg, Dominik Macak, Tomislav Maricic, Linda Dombrowski, Maria Schörnig, Konstantinos Anastassiadis, Oliver Baker, Ronald Naumann, Ekaterina Khrameeva, Anna Vanushkina, Elena Stekolschikova, Ilia Kurochkin, Randall Mazzarino, Nathan Duval, Dmitri Zubkov, Patrick Giavalisco, Terry G. Wilkinson, David Patterson, Philipp Khaitovich, Svante Pääbo

## Abstract

We analyze the metabolomes of humans, chimpanzees and macaques in muscle, kidney and three different regions of the brain. Whereas several compounds in amino acid metabolism occur at either higher or lower concentrations in humans than in the other primates, metabolites in oxidative phosphorylation and purine biosynthesis are consistently present in lower concentrations in the brains of humans. In particular, metabolites downstream of adenylosuccinate lyase, which catalyzes two reactions in purine synthesis, occur at lower concentrations in humans. This enzyme carries an amino acid substitution that is present in all humans today but absent in Neandertals. By introducing the modern human substitution into the genomes of mice, as well as the ancestral, Neandertal-like substitution into the genomes of human cells, we show that this amino acid substitution is responsible for much or all of the reduction of *de novo* synthesis of purines in humans.

## Introduction

Modern humans differ dramatically from their closest evolutionary relatives in a number of ways. Most strikingly, they have developed rapidly changing and complex cultures that have allowed them to become more numerous and widespread than any other primates and closely related hominins such as Neandertals. This unique historical development has at least to some extent biological roots. However, although large numbers of traits have been identified as being unique to humans or been suggested to be so (*e*.*g*. (Tomasello, 2019); (Varki and Altheide, 2005)), it has proven difficult to identify the genetic and biological underpinnings of such traits. One reason may be that many of them are genetically complex (Phillips, 2008) and represent differences in degree rather than absolute differences between humans and other groups. In particular, it is unclear if any single nucleotide substitution that occurs in all or almost all humans today, but is not seen among the closest evolutionary relatives of present-day humans, the Neandertals and Denisovans (Pääbo, 2014), has biological consequences.

To identify traits that may have comparatively simple genetic architectures yet influence the biology of modern humans, we focus here on metabolic differences. By analyzing the metabolomes of muscle, kidney and three different regions of the brain from humans, chimpanzees and macaques, we find that many aspects of amino acid metabolism differ between humans and the other two primates in all tissues analyzed. Among metabolic pathways, oxidative phosphorylation is less active in the brains of humans than the other primates and purine biosynthesis is less active in the human brain as well as in other tissues. In purine biosynthesis, we find that metabolites downstream of the enzyme adenylosuccinate lyase [ADSL (EC 4.3.2.2)] occur at lower concentrations in humans than in the other primates. ADSL carries an amino acid substitution that is unique to modern humans relative to apes and Neandertals and Denisovans (Castellano et al., 2014) and has been shown to affect the stability of the enzyme (Van Laer et al., 2018). By introducing the modern human-like substitution in the genome of mice, and the ancestral, Neandertal-like substitution in the genomes of human cells, we show that this substitution is responsible for much or all of this metabolic change in present-day humans.

## Results

### Metabolite changes unique to humans

To identify metabolites that have changed their concentration in humans relative to monkeys and apes, we measured compound concentrations in prefrontal cortex, primary visual cortex, cerebellum, skeletal muscle and kidney in four humans, four chimpanzees, and four macaques using mass spectrometry coupled with capillary electrophoresis (CE-MS), a technique suitable for detection of small hydrophilic compounds. The number of metabolites annotated in the five tissues varied between 166 and 197, 160 and 209, and 145 and 192 in the three species, respectively (Supplementary Table 1).

In each tissue, we identified metabolites that did not differ significantly in concentrations between the macaques and the chimpanzees, but differed significantly, and in the same direction, between humans and macaques as well as between humans and chimpanzees. Whereas we find no such metabolites in skeletal muscle and kidney, we find metabolites that differ in their concentration in humans relative to the other two primates in two of the three parts of the brain analyzed.

In cerebellum, 22 metabolites have higher concentrations in humans whereas no metabolites have lower concentrations (Supplementary Table 2). Eighteen of the 22 metabolites are amino acids. In prefrontal cortex we detected five metabolites with lower concentrations in humans and none with higher concentrations (Supplementary Table 2). Three of the five metabolites are purines (inosine monophosphate (IMP), guanosine monophosphate (GMP), adenosine monophosphate (AMP)) and the other two are NAD^+^ and UDP-N-acetylglucosamine.

### Metabolic pathways

To identify metabolic pathways that may be more or less active in humans than in other primates, we linked metabolites with higher or lower concentrations in humans compared to both chimpanzees and macaques (even if not significantly so) to genes and metabolic pathways using the Kyoto Encyclopedia of Genes and Genomes (KEGG) database. Supplementary Tables 3a and b give pathways that contain more genes than expected (p<0.01) that are associated with metabolites present in either higher or lower concentrations in each of the five tissues.

Between 8 and 12 pathways are associated with higher metabolite concentrations in the five human tissues (Supplementary Table 3a). All eight pathways identified in the prefrontal cortex involve amino acid metabolism, eight out of nine pathways in the visual cortex and nine out of ten in the cerebellum also involve amino acids. In muscle and kidney, seven and five of the eleven and twelve pathways identified, respectively, involve amino acid metabolism. Thus, several aspects of amino acid metabolism seem to be increased in humans relative to other primates.

Between four and eleven pathways are associated with lower metabolite concentrations in the five tissues (Supplementary Table 3b). Amino acid and peptide metabolism make up one to five of these pathways in the different tissues, suggesting that amino acid metabolism has changed in several ways in humans relative to the other primates. Hence, metabolites in these pathways are present in increased as well as decreased concentrations, often in the same pathways. For example, significant numbers of metabolites in arginine and proline metabolism are present in higher concentrations while other metabolites in the same pathways are present in lower concentrations, sometimes in the same tissues, suggesting that the flux through these pathways has changed.

In contrast, metabolites in two pathways are consistently present at lower rather than higher concentrations in humans. One of these pathways is oxidative phosphorylation, which show decreased metabolite concentrations in all three brain regions analyzed. Because oxidative phosphorylation is not affected in muscle and kidneys, it seems that the human brain differs from ape brains in that oxidative phosphorylation in mitochondria is less active.

The second pathway that stands out is purine biosynthesis where a number of metabolites are present at lower levels in all five tissues analyzed. Thus, purine biosynthesis is decreased in humans relative to apes in the brain as well as in other organs. If we focus on a more restrictive definition of purine biosynthesis (Marie et al., 2004), a significant decrease is seen only in the three brain regions and not in muscle and kidney (Figure 1a). Furthermore, three of the five metabolites, IMP, GMP and AMP, which individually show significantly human-specific concentration decreases, are products of *de novo* purine biosynthesis. Thus, purine biosynthesis stands out as down-regulated in humans, particularly in the brain.

**Figure 1.**
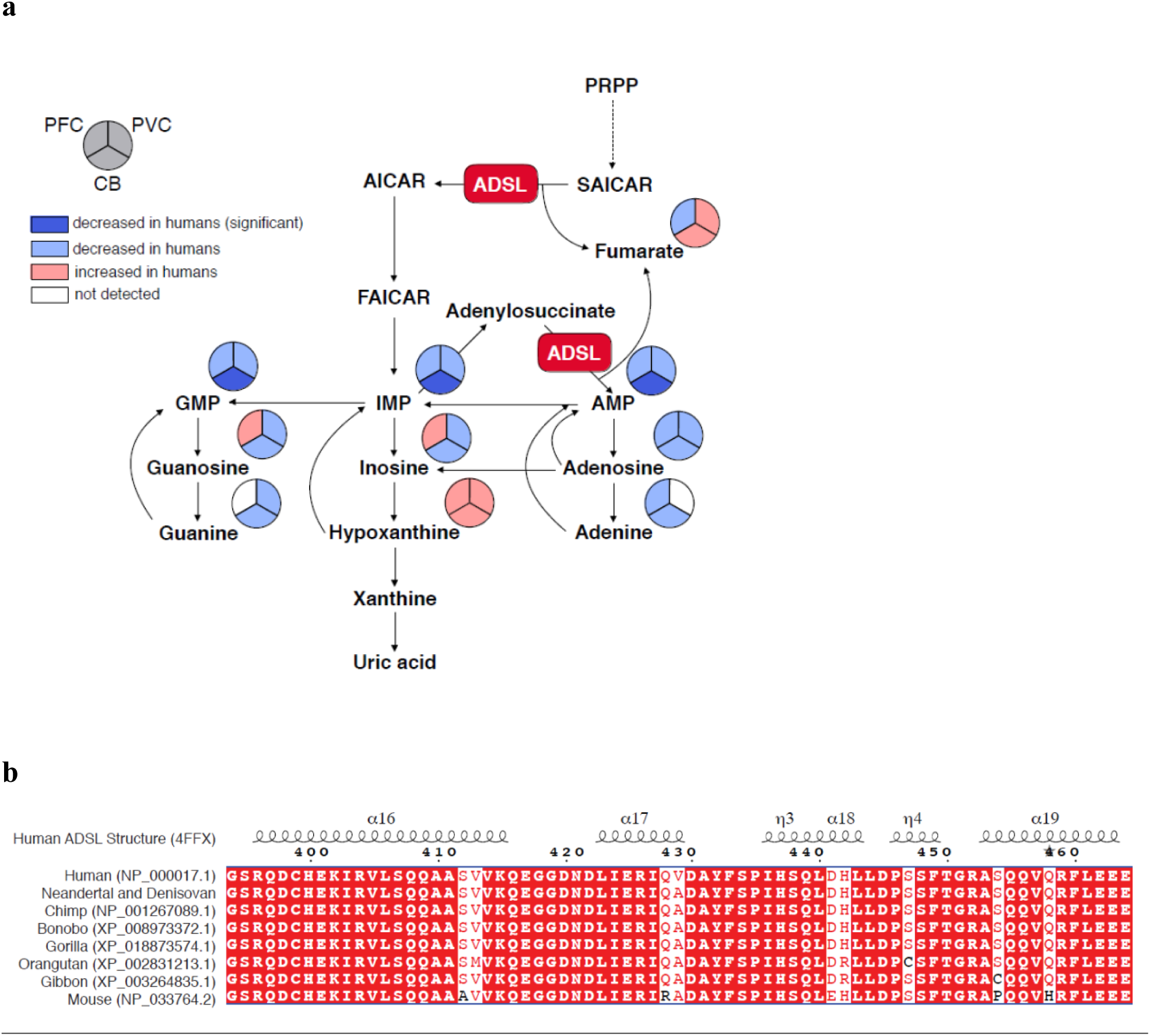
Human-specific differences of metabolite levels in purine biosynthesis pathway and ADSL protein sequence. **a**. Purine biosynthesis in humans relative to apes. Pathway sketch illustrating the central role of ADSL in purine biosynthesis and showing the human-specific changes when compared to chimpanzees and macaques in metabolite concentrations in three brain regions as detected by CE-MS. PFC – prefrontal cortex, PVC – primary visual cortex, CB – cerebellum. **b**. Partial amino acid sequences of ADSL. Accession numbers and amino acid sequences are from NCBI. X-ray structural determination of ADSL (4FFX) is from (Ray et al., 2012).

### Adenylosuccinate lyase (ADSL) in modern humans

The enzyme adenylosuccinate lyase (ADSL) catalyzes two reactions in *de novo* purine biosynthesis (Ciardo et al., 2001; Jaeken and Van Den Berghe, 1984). First, it cleaves succinylaminoimidazole carboxamide ribotide (SAICAR) into aminoimidazole carboxamide ribotide (AICAR) and fumarate. Second, it cleaves adenylosuccinate (S-AMP) into adenosine monophosphate (AMP) and fumarate (Figure 1a). The compounds AMP, IMP, and GMP that are reduced in the three human brain regions analyzed are situated downstream of ADSL, consistent with the hypothesis that a change in ADSL may have caused the reduction in purine biosynthesis.

ADSL (Ariyananda et al., 2009) is a homotetrameric complex where three monomers contribute to each of the four active sites (Brosius and Colman, 2002). The gene encoding ADSL in humans is located on chromosome 22q13.1–13.2 and is one of only about a hundred protein-coding genes that carry substitutions inferred to change amino acids that are fixed or almost fixed in present-day humans but occur in their ancestral, ape-like state in the genomes of the closest extinct relatives of modern humans, Neandertals and Denisovans (Pääbo, 2014). The substitution in *ADSL* in modern humans is an alanine to valine substitution at position 429 (A429V). Neandertals, Denisovans, all other primates and most mammals carry an alanine residue at this position (Figure 1b). The substitution is located in a protein domain that forms part of the substrate channel over the active site of the enzyme. Amino acid substitutions close to this position cause lowered enzymatic activity and/or lower enzyme stability and result in symptoms that include psychomotor retardation, autism, and epilepsy (Jurecka et al., 2015; Spiegel et al., 2006; Zikánová et al., 2010) as well as alterations in brain structure as seen by magnetic resonance imaging (Jurecka et al., 2012). This raises the possibility that the amino acid substitution at position 429 may have changed the activity of ADSL in modern humans after their separation from the lineage leading to Neandertals and Denisovans.

### A mouse humanized for *Adsl*

To investigate how the A429V substitution may affect purine biosynthesis, we introduced this mutation into the *Adsl* gene of a C57BL/6N mouse by injection of the relevant oligonucleotide and CRISPR-*Cas9* into the male pronucleus of fertilized oocytes. Adjacent to the amino acid of interest, at position 428, mice carry an arginine residue whereas primates carry a glutamine residue (Figure 1b). To avoid possible effects of the rodent-specific arginine residue at position 428 on the function of the amino acid at position 429, we introduced a nucleotide substitution resulting in an R428Q substitution in addition to the V429A substitution in the *Adsl* gene. The two mutations segregate in Mendelian ratios in the mice, and animals heterozygous and homozygous for the two substitutions show no overt phenotypic difference to their wild type littermates.

### ADSL activity in the humanized mice

We analyzed the enzymatic activities of ADSL in tissue extracts prepared from mice homozygous for the human-like mutations and their wild-type littermates in cerebellum, cerebral cortex, heart, kidney, liver, lung, muscle, spleen and testis in 12-week-old mice and (except testis) in one-week-old mice by measuring the conversion of adenylosuccinate to AMP.

In the wild-type adults, mean ADSL activity varied from 1.95 nmole/min/μg in liver to 24.4 nmole/min/μg in muscle (Figure 2). In the pups, mean enzymatic activity was 5.4% to 57% higher in most tissues and largely paralleled those of the adults. Two exceptions were cerebellum and muscle, where ADSL activity was 41% and 11% lower in the pups than in the adults, respectively.

**Figure 2.**
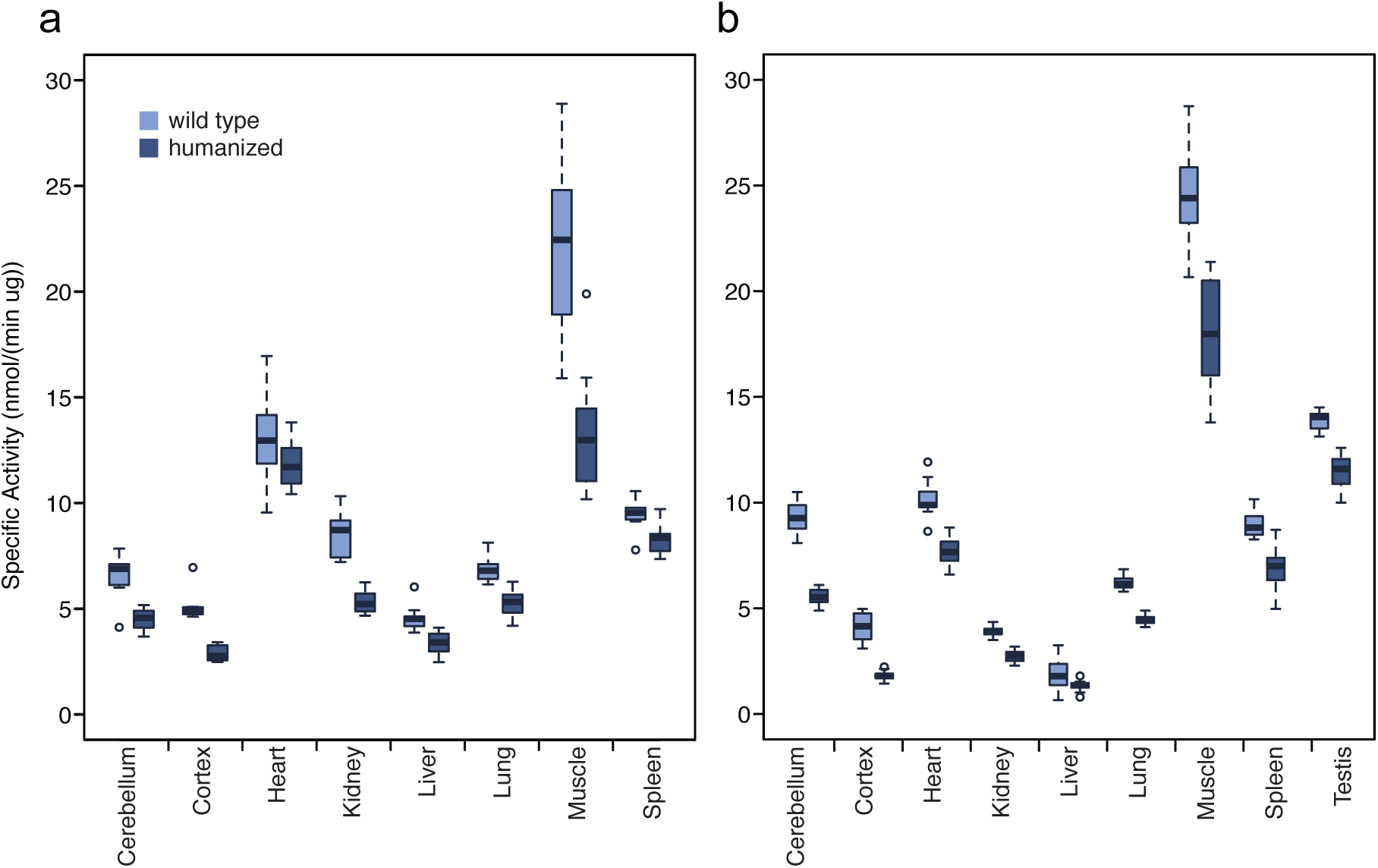
ADSL activity in tissues from humanized and wild type mice. **a**. One-week-old mice. **b**. Twelve-week-old mice. Conversion of adenylosuccinate to AMP was monitored by measuring λ 282 nm absorbance over 20 minutes (see Methods).

In the humanized mice, ADSL activity was reduced by 8.7 to 56% in all organs in both adults and pups (p<0.05) (Figure 2). The relative reduction of the activity in the humanized relative to the wild-type mice is higher in cerebral cortex (43% in pups and 56% in adults) than in the other tissues (8.7% - 40%). Thus, young as well as adult mice that carry an ADSL enzyme humanized at positions 428 and 429 have lower ADSL activity in most or all tissues.

### The metabolome of humanized *Adsl* mice

We next analyzed the metabolomes of the nine tissues from 9-12 adult homozygous humanized mice and their wild-type littermates by gas chromatography coupled with mass spectrometry (GC-MS). We similarly analyzed eight tissues (no testis) from 10-12 one-week-old pups (Supplementary Table 4). The number of metabolites detected varied between 176 and 273 in the adult mice and between 310 and 347 in the young mice. Among those, an average of 280 (median = 273) were detected in at least 50% of the adult and young individuals in each of the nine tissues (Supplementary Table 5a,b). A principle component analysis using the concentrations of all metabolites detected revealed one to two outlier samples per tissue (Supplementary Table 4). These were excluded from further analyses. Including these samples in the analyses did not qualitatively affect results.

Among the organs analyzed, only the brain showed significant differences in metabolite concentrations between the wild type and humanized mice. Specifically, 36 metabolites showed significant concentration differences in cerebellum of 12-week-old mice (permutations, p = 0.036) and 45 metabolites showed marginally significant differences in the cerebral cortex of one-week-old mice (permutations, p = 0.081, Supplementary Table 6). Thus, the metabolic effects of the two mutations introduced in ADSL are particularly pronounced in the central nervous system.

The concentration differences detected in cerebellum of the 12-week-old mice correlated with differences observed in cerebral cortex, even though differences in cortex did not pass the significance cut-off in the permutation test (Pearson correlation, r = 0.84, p < 0.0001, n = 29; Supplementary Fig. 1a). Similarly, concentration differences detected in cerebral cortex in one-week-old mice correlated with differences observed in cerebellum in the same mice (Pearson correlation, r = 0.77, p < 0.0001, n = 45; Supplementary Fig. 1b). The concentration differences between wild type and humanized mice furthermore correlated between the one-week-old and 12-week-old mice both in cortex and in cerebellum (Pearson correlation, r = 0.49 and r = 0.52, p < 0.008, n = 28 and n = 31, respectively; Supplementary Fig. 1e, f). By contrast, the correlation between metabolic differences detected in brain and the other tissues was weaker (p < 10^−8^ for brain tissues and p > 10^−4^ for other tissues) (p < 10^−8^; Supplementary Fig. 1c,d). Thus, in the humanized mouse, the effects of the substitutions in ADSL are seen in the cerebral cortex and cerebellum in both young and adult animals.

### Purine biosynthesis in the humanized mice

In the 12-week-old mice, we detected six metabolites within the purine biosynthesis pathway. In the humanized mice, four of these metabolites had lower concentrations than in the wild-type mice in the cerebral cortex and all of them had lower concentrations in the cerebellum (binomial test for cortex and cerebellum, p = 0.04) (Figure 3).

**Figure 3.**
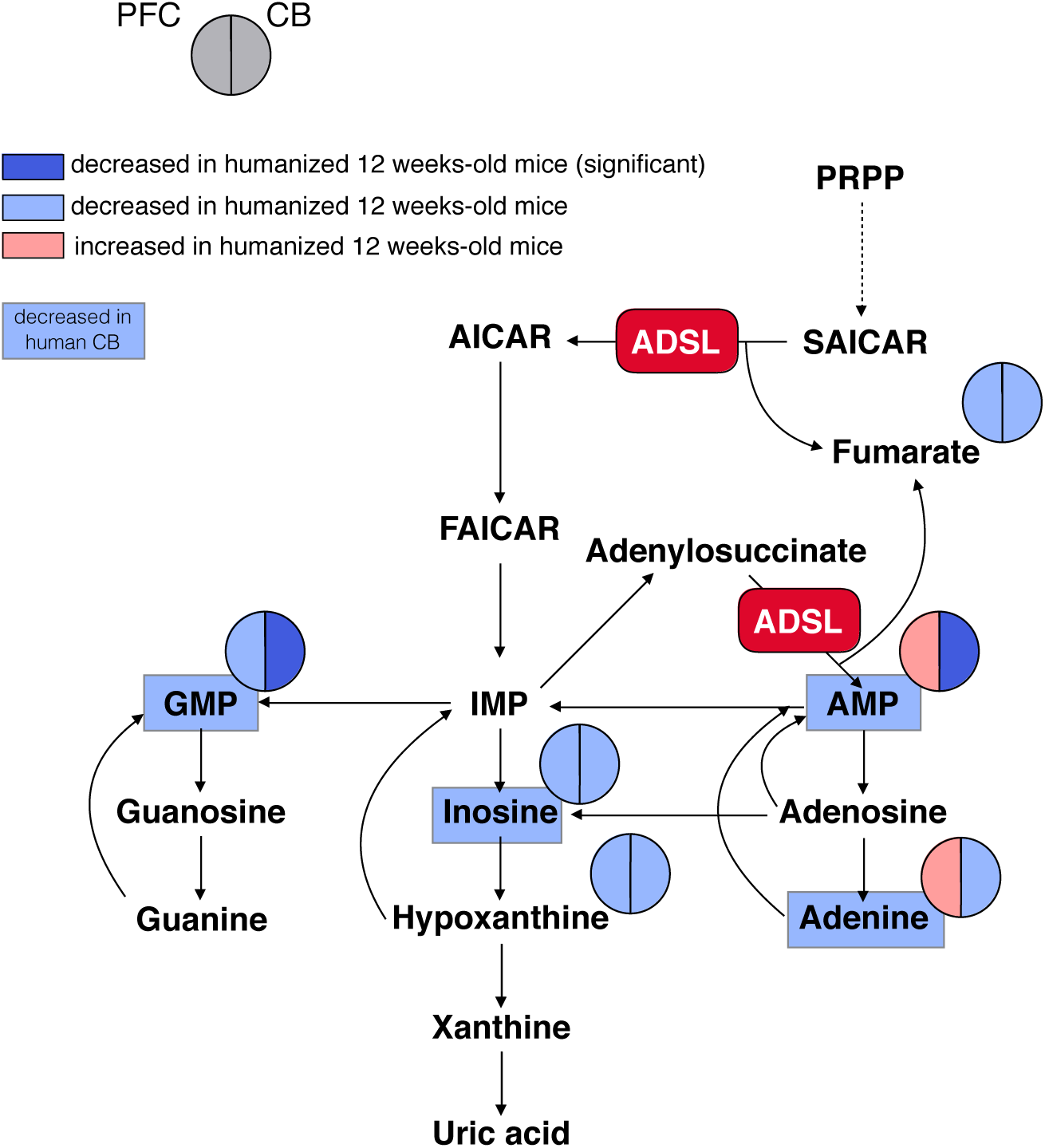
Purine biosynthesis in adult mice “humanized” for ADSL. Pathway sketch showing changes in metabolite concentrations as detected by CE-MS in mice carrying the V429A and R428Q substitutions in the ADSL protein. PFC – prefrontal cortex, CB – cerebellum. Metabolites that are also present in lower concentrations in human cerebellum than in chimpanzee are on blue background.

The six metabolites detected in 12-week-old mice were also detected in the primate brains. Four of these had lower concentrations in the human brain, including AMP and GMP in prefrontal cortex, the visual cortex, and in cerebellum (binomial test, p = 0.04). Furthermore, four of the six metabolites with lower concentrations in cerebellum in the humanized mice had lower concentrations in human cerebellum compared to other primates: AMP, GMP, inosine, and adenine (Figure 3). In the one-week-old mice, we detected 7 metabolites in the purine biosynthesis pathway. In the humanized mice, five of these had lower concentration in the cerebral cortex and three of these had lower concentration in the cerebellum when compared to their wild-type litter mates (Supplementary Fig. 2).

Thus, several changes in concentrations of compounds in purine biosynthesis seen in the humanized mice recapitulate differences seen when the metabolomes of human brains are compared to the brains of chimpanzees and macaques.

### Activity and stability of humanized mouse ADSL

To investigate how the humanized form of the mouse ADSL enzyme may influence purine biosynthesis, we synthesized mouse wild-type (wt) ADSL and mouse A429V ADSL and inserted them in expression vectors that include N-terminal polyhistidine tags (pET-14b vector) (Lee and Colman, 2007). We analyzed the conversion of SAICAR to AICAR, and the conversion of SAMP to AMP, in the presence of excess substrate by measuring the rate of production of AICAR and AMP (Figure 4a) and found no differences in the kinetics of either reaction between wt and A429V ADSL (t-test, p>0.05).

**Figure 4.**
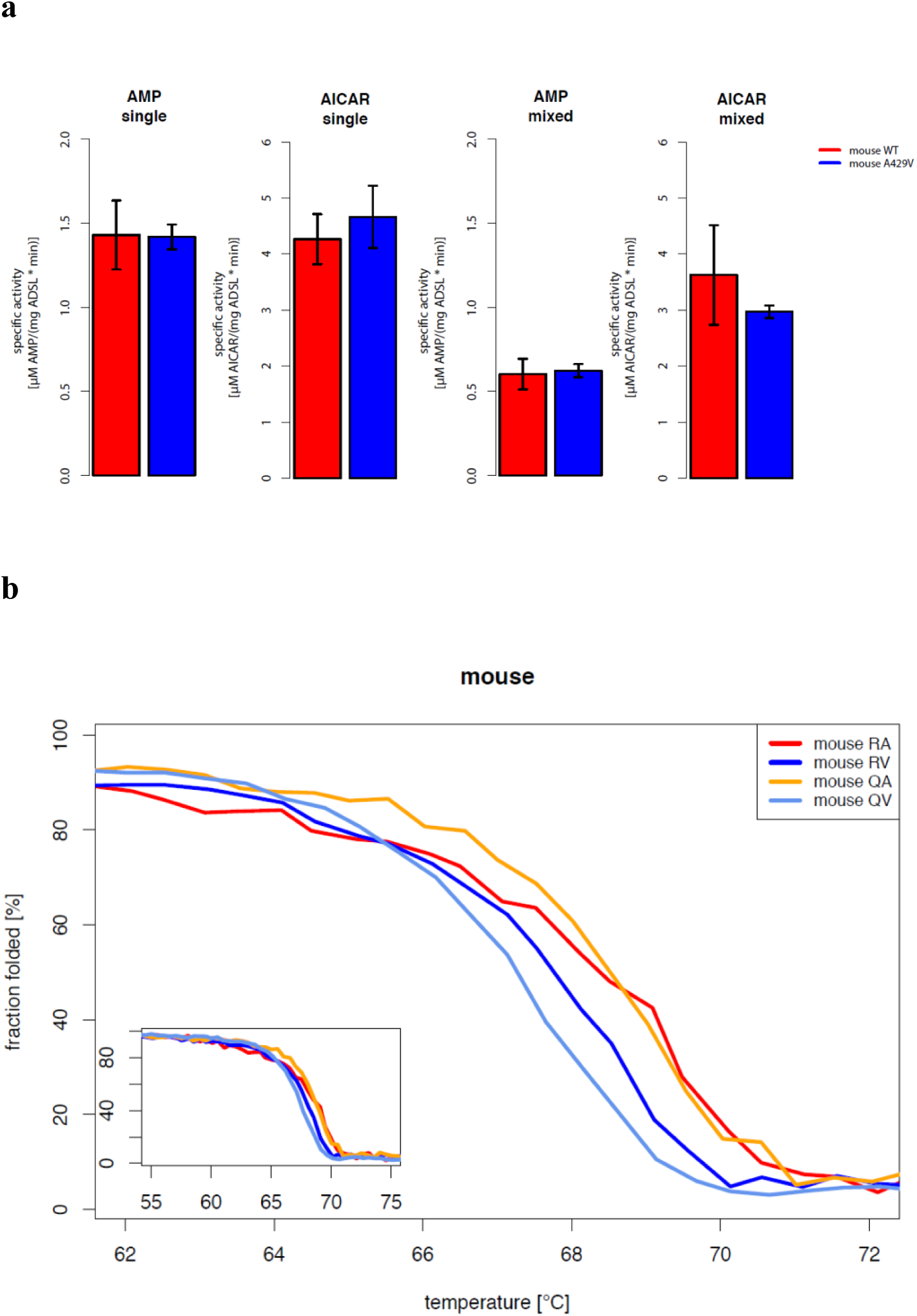
Characterization of mouse ADSL forms. **a**. Enzyme kinetics tested in excess of a substrate for the products (AICAR and AMP, see text above the plots) with one substrate (single) or with both substrates (mixed) in the reaction mix. Specific activity of wild type and humanized A429V versions do not differ (P>0.05, t-test). **b**. Protein melting measured by CD at 222nm for mouse WT (RA), mouse RV, mouse QA and mouse QV ADSL versions. The RV (p=0.10, t-test) and QV versions (p=0.05, t-test) are less stable.

We next investigated if the A429V substitution, and/or the R428Q substitution that was introduced into the humanized mice together with the A429V substitution, affect the conformational stability of the protein by constructing expression vectors that carry wild-type murine ADSL, ADSL with only the A429V substitution, ADSL with only an R428Q substitution, and ADSL with the A429V substitution and the adjacent R428Q substitution. We then compared the secondary structure stability of all four protein variants by measuring the circular dichroism spectra of the purified proteins at 222 nm while heating them from 55 °C to 80 °C at a rate of one degree per minute (Figure 4b). Mouse ADSL protein with alanine at position 429 was more stable (50% folded at 68.3 °C, +/-0.3 °C, n=4) than mouse ADSL protein with valine at position 429 (50% folded at 67.3 °C +/-0.3 °C, n=4). The effect on the protein stability of the A429V substitution was not influenced by the presence or absence of the adjacent R428Q substitution at position 428. In summary, the A429V substitution does not affect the kinetics of the murine ADSL enzyme. However, it decreases the secondary structure stability of the protein.

### Characterization of human and Neandertal ADSL

Previous work has shown that the modern human version of ADSL is less stable than the ancestral, Neandertal-like form *in vitro* (Van Laer et al., 2018). To ensure that the two forms of the ADSL protein which differ only at position 429 in the protein are both enzymatically active in living cells we show that they both are able to rescue Chinese hamster ovary cells that lack ADSL activity (*AdeI* cells, (Vliet et al., 2011)) (Supplementary Fig. 3a). We then expressed the modern human and Neandertal versions of ADSL in *E. coli* and isolated the two proteins by N-terminal His-tags (Lee and Colman, 2007). In agreement with previous results (Van Laer et al., 2018), circular dichroism spectra of both protein versions exhibited predominantly alpha-helical characteristics (Supplementary Fig. 3b). To investigate if the enzymatic activities of the human and Neandertal versions of ADSL differ, we tested their ability to convert SAICAR to AICAR, and SAMP to AMP, as described above. No differences in specific activity between the two enzymes were detected using either substrate (SAMP, t-test, p=0.41 or SAICAR, t-test, p=0.81, Figure 5a).

**Figure 5.**
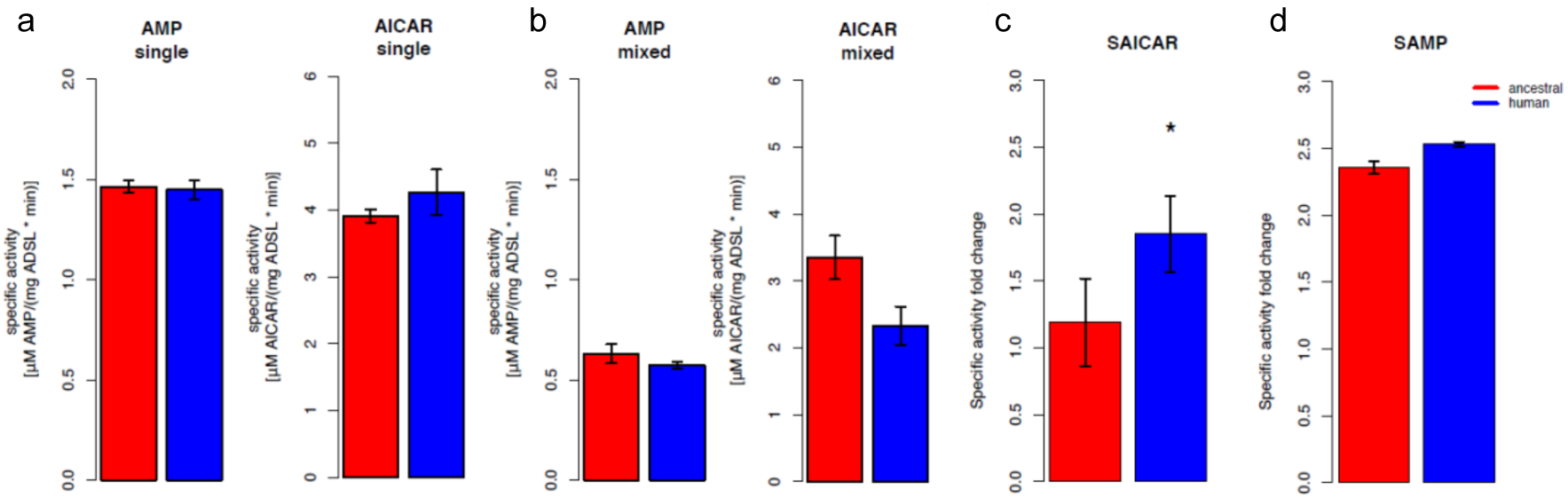
Specific activity and cooperativity of human and ancestral ADSL versions. Specific activity of SAMP to AMP and SAICAR to AICAR conversions for the Neandertal-like (red) and modern human (blue) forms of ADSL when incubated with each substrate separately **a**, or with both substrates (“mixed”) **b** (P>0.05, t-test). **(c** and **d)** Changes in enzyme cooperativity between ancestral and human versions given as the ratios of activities in presence of single substrates to both substrates. The human version show a higher reduction of activity for SAICAR to AICAR conversion in the presence of both substrates than the ancestral version (t-test, p= 0.01) **c**, which is not seen for SAMP to AMP conversion **d** (t-test, p= 0.49). Bars represent average values over 3 repetitions, error bars represent standard errors.

To investigate if cooperativity between the two activities of ADSL may be affected by the A429V substitution, we incubated the two forms of the enzyme with equimolar amounts of SAMP and SAICAR and calculated relative values of ADSL protein activity when presented with a single substrate (Figure 5a) or an equimolar mixture of SAMP and SAICAR (Figure 5b). The conversion of SAMP to AMP is reduced approximately to the same extent (∼2.4-fold, t-test, p=0.49) for the modern human and Neandertal-like forms of the enzyme when both substrates are present (Figure 5d). In contrast, the conversion of SAICAR to AICAR is reduced about 1.8-fold for the modern human form in the presence of both substrates (p=0.026) while the ancestral form of the enzyme shows no reduction (Figure 5c) (p=0.44). This suggests that the modern human enzyme produces less AICAR than the ancestral enzyme under physiological conditions and may warrant further investigation.

### Stability of modern human and ancestral ADSL

To investigate the effects of the A429V substitution in the human protein, we compared the stability of the secondary structure of the protein as described above. The temperature at which 50% of the protein is inferred to be folded is 67.0 °C (+/-SEM=0.10 °C) for the modern human form of the protein and 68.0 °C (+/-SEM=0.2 °C) for the Neandertal-like form of the protein (Figure 6a). Thus, as previously shown (Van Laer et al., 2018), modern human ADSL is less stable than ancestral ADSL.

**Figure 6.**
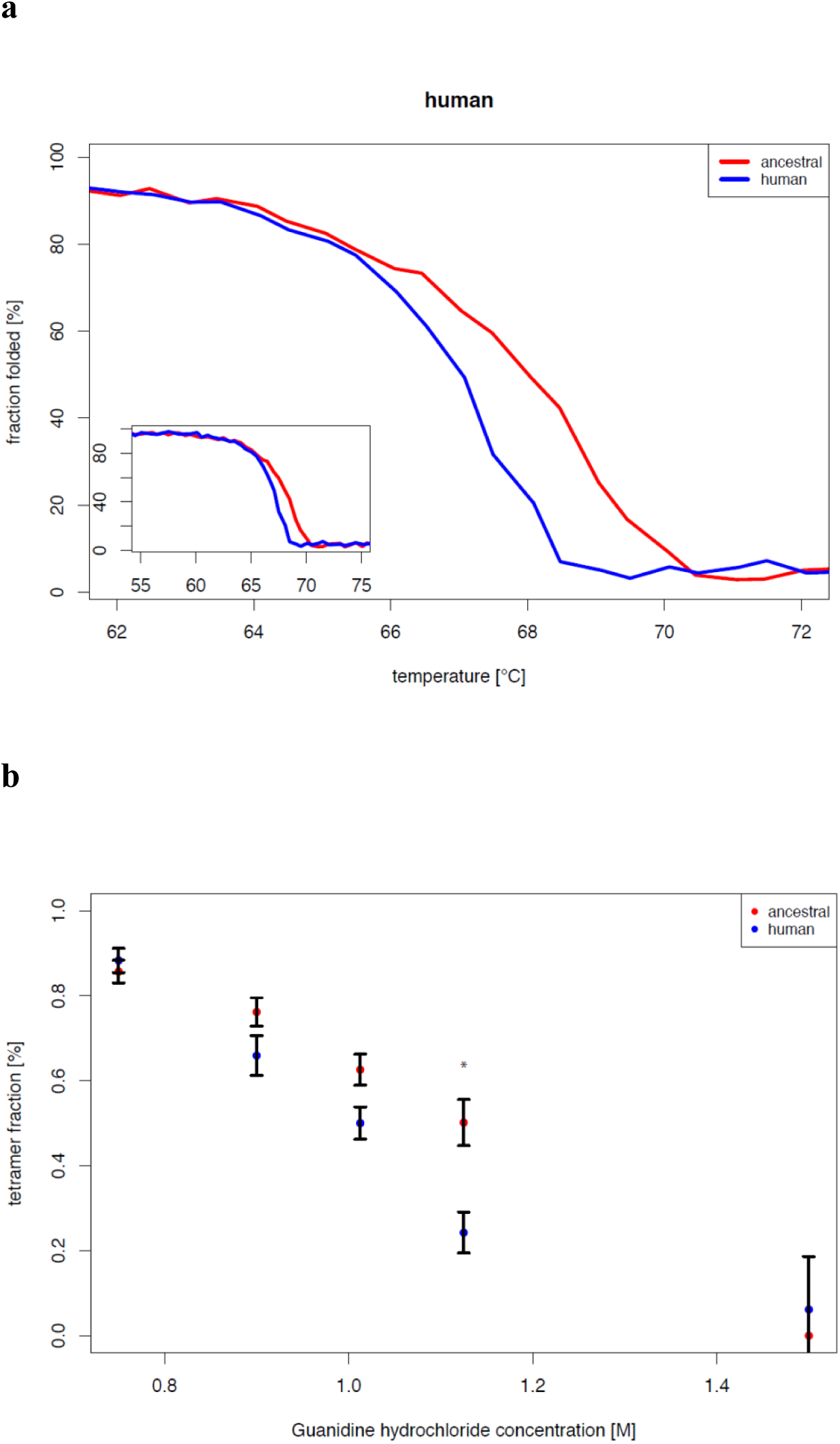
Stability of modern human and Neandertal-like ADSL. **a**. Thermal denaturation as observed as CD at 222nm for 55°C-75°C (plot insert) and enlarged for 62-72°C (bigger panel) indicate lower stability of the modern human ADSL at the denaturation midpoint (t-test, p=0.001). **b**. Chemical denaturation observed by native gel electrophoresis (p=0.024 for 1.125M Gdn-Hcl concentration, t-test with Bonferroni correction). Plot represents average amount of tetrameric protein over 5 repetitions. Error bars represent standard errors.

To investigate the effect of the difference in secondary structure stability on the stability of ADSL quaternary structures, we exposed the proteins to 0.75M, 0.90M, 1.0125M, 1.125M and 1.5M guanidine hydrochloride (GdnHCl) overnight at room temperature and analyzed the proteins by native gel electrophoresis. Bands corresponding to the predicted size of the ADSL tetramer (∼223 kDa) and monomer (∼56 kDa) were observed and the relative proportion of tetrameric protein was estimated. Figure 6b shows that at 1.0125 M GdnHCl about 50% of the modern human version of ADSL was present as tetramers whereas for the ancestral form this was the case at 1.125 M GdnHCl. Thus, both the secondary and quaternary structures of the ancestral form of ADSL are more stable than for the modern human form.

### Ancestralized ADSL in human cells

To investigate how the A429V substitution affects the metabolome of living human cells, we used CRISPR-*Cas9* to introduce the nucleotide substitution in the *ADSL* gene that reverts the valine at position 429 to the ancestral alanine residue in human 409B2 cells (Riken BioResource Center). We isolated three independent cell lines and we verified that the intended nucleotide substitution had occurred in each of them by sequencing a segment of the *ADSL* gene and excluded deletions of the target site by quantitative PCR (not shown). We also isolated six independent lines that had been subjected to the CRISPR-*Cas9* procedure but did not exhibit any mutation in the sequenced DNA segment. We expanded 10 separate cultures of the three edited cell lines and 19 cultures of the control lines. Each of these cultures were analyzed by LC-MS in positive and negative modes.

A total of 10,673 metabolites were detected. Among these were twelve metabolites from the *de novo* purine biosynthesis pathway and three of these (aminoimidazole ribonucleotide (AIR), hypoxanthine, guanine) were present at significantly lower amounts in the wild type versus the ancestralized cells (p<0.01). Strikingly, out of nine detected metabolites downstream of ADSL, all have lower concentrations in the wild type cells (binom. test, p = 0.002, Figure 7). Thus, in human cells, the ancestral form of ADSL supports a higher level of purine biosynthesis than the modern human form.

**Figure 7.**
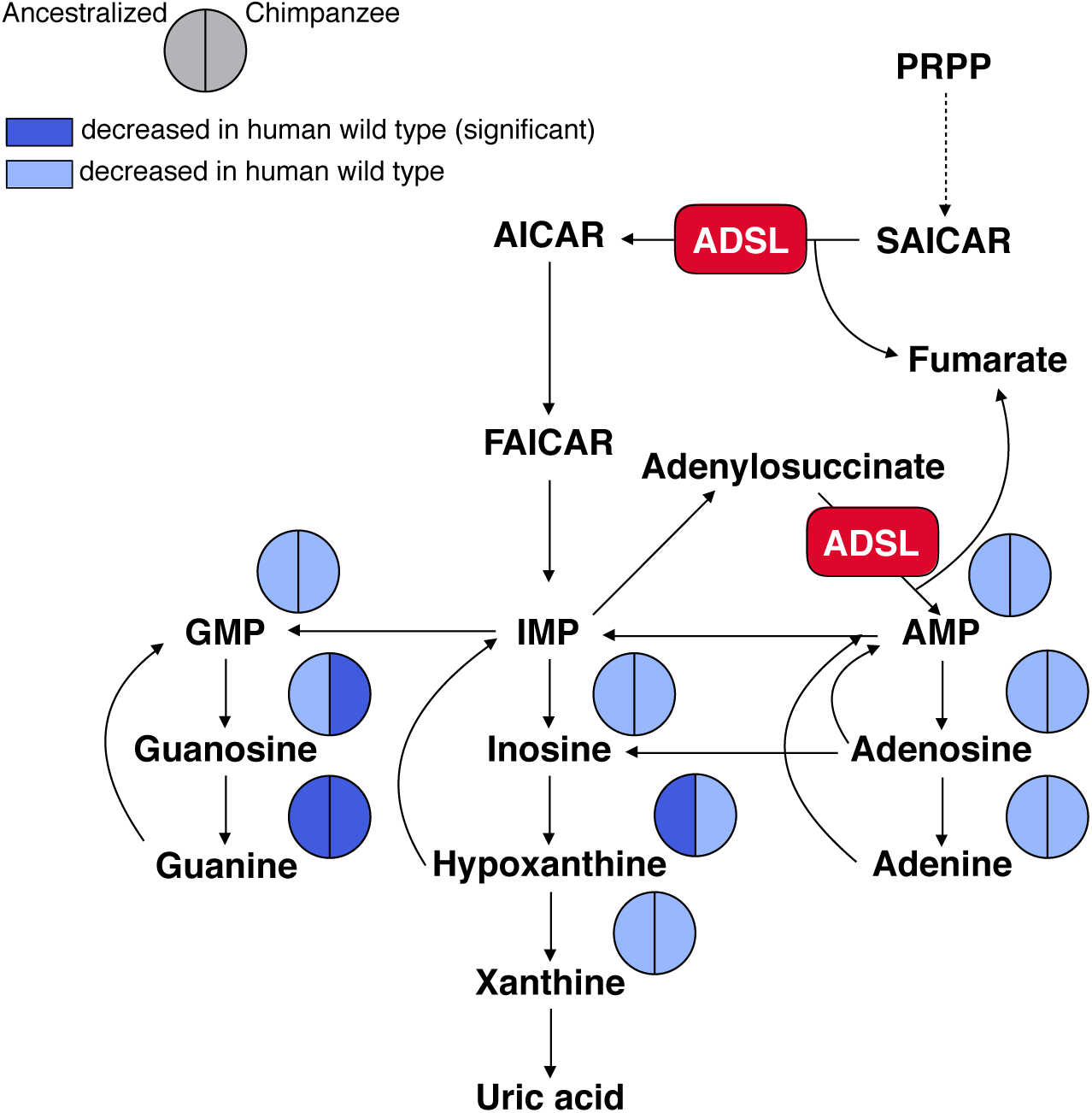
Purine biosynthesis in human cells carrying the V429A substitutions and chimpanzee cells compared to wild type cells.

### Comparison to chimpanzee cells

To compare these results to purine biosynthesis in chimpanzees, we similarly analyzed nine cultures from three different chimpanzee pluripotent cell lines. Out of 1,286 metabolites that differ significantly (p<0.01) between the human and chimpanzee wild type cells, three purine biosynthesis metabolites (AIR, guanosine, guanine) are at higher concentrations in the chimpanzee cells. The same nine metabolites as in the comparison to the ancestralized human cells were detected in the chimpanzee cells. Similar to the ancestralized human cells, they are present in lower concentrations in the human than in the chimpanzee cells (binom. test, p = 0.002, Figure 7). Thus, reversal of the A429V substitution in ADSL in human cells results in an increase in purine biosynthesis similar to what is observed when chimpanzee cells are compared to human cells.

### AMPK dephosphorylation

One way in which lower steady state levels of metabolites downstream of ADSL in purine biosynthesis may affect cells is through activation of the AMP-activated protein kinase (AMPK). This serine/threonine protein kinase complex is composed of a catalytic α-subunit, a scaffolding β-subunit and a regulatory γ-subunit. Of particular relevance may be phosphorylation of the threonine at position 172 in the α-subunit [pAMPKα(Thr172)], which is enhanced when AMP is bound by the γ-subunit (Davies et al., 1995) (Hawley et al., 1995). AICAR, a direct product of ADSL and an intermediate in the generation of IMP and AMP, can mimic AMP in stimulating AMPK activity (Daignan-Fornier and Pinson, 2012; Kim et al., 2016; Mihaylova and Shaw, 2011). As SAICAR to AICAR catalysis (Figure 5c) as well as AMP levels (Figure 7) are reduced in human cells, phosphorylation of AMPKα(Thr172) may be reduced, and its activity decreased.

To test if this is the case, we measured the levels of phosphorylated AMPKα(Thr172) in two wild type human and two ancestralized cell lines by immunoblotting with antibodies specific to pAMPKα(Thr172) and to total AMPKα. The levels of phosphorylated AMPKα(Thr172) were 28.5% higher in the ancestralized cells than in the wild type human cells (p=0.04) (Figure 8a) and the ratio of phosphorylated AMPKα to total AMPKα was 10.3% higher in the ancestralized cells (p=0.10), consistent with the hypothesis that higher levels of AICAR and AMP in the ancestralized human cells may increase the phosphorylation of AMPK. Next, we compared the pAMPKα(Thr172) levels in three human and three chimpanzee cell lines. The levels of phosphorylated AMPKα(Thr172) were 59.3% higher in the chimpanzee than human the cells (p=0.03) (Figure 8b), while the ratio of phosphorylated AMPKα to total AMPKα was 47.9% higher in the chimpanzee cells (p=0.002). Thus, ancestralization of position 429 in ADSL in human cells reduces purine biosynthesis and phosphorylation of threonine at position 172 in the α-subunit of AMPK and both effects are qualitatively similar to differences seen when chimpanzee and human cells.

**Figure 8.**
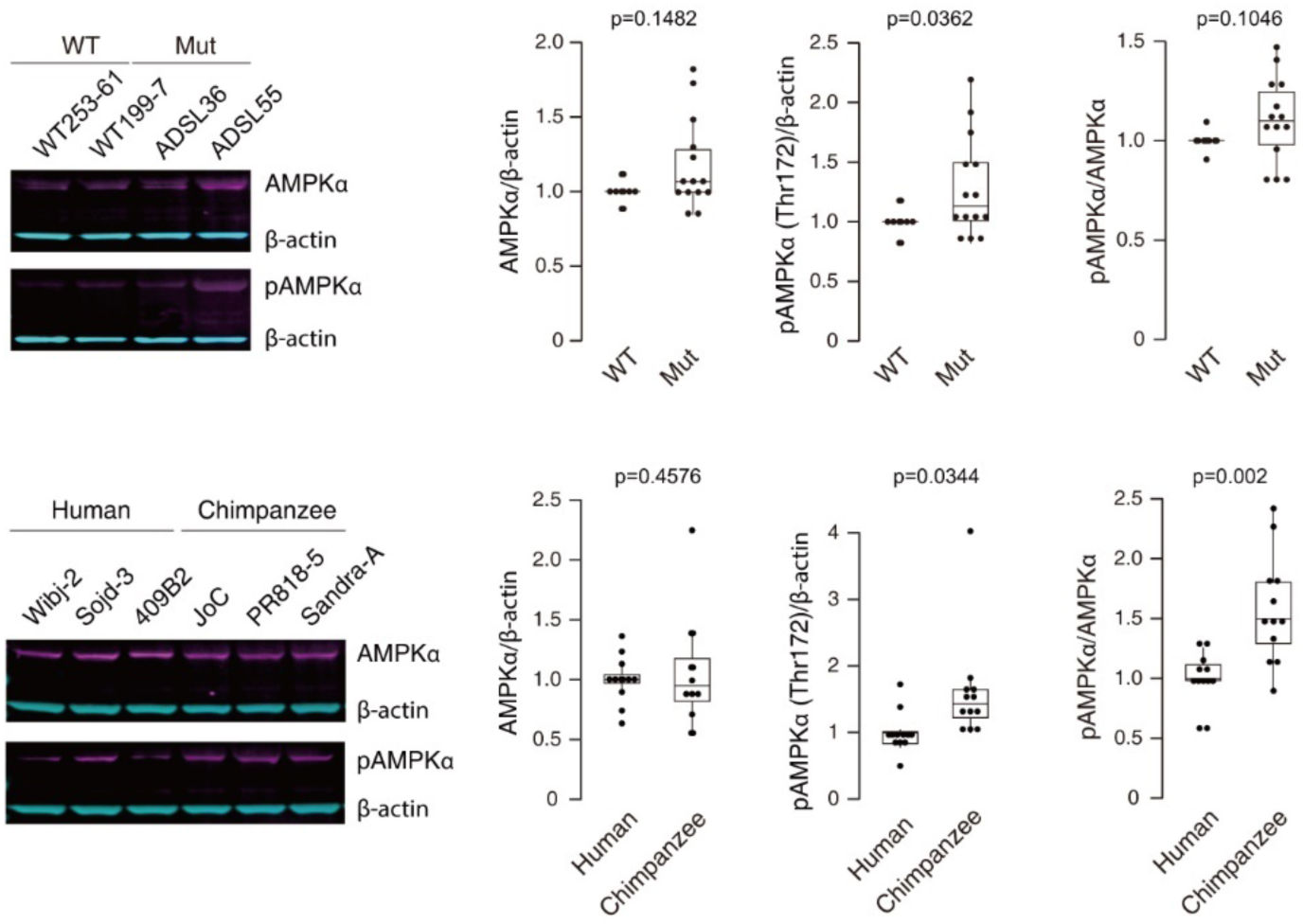
Total AMPK and phosphorylated AMPKα(Thr172) in ancestralized human and chimpanzee cells. **a**. Results for two wild type (WT) cellular clones (253-61, 199-7) and two cellular clones (ADSL36, ADSL55) carrying the V429A substitutions (Mut) are shown, as well as the ratios of the quantitation of the proteins. **b**. Results for three human and three chimpanzee cell lines.

## Discussion

The ancestors of modern humans diverged from their closest evolutionary relatives, Neandertals and Denisovans, on the order of 600,000 years ago (Prüfer et al., 2014). About ten times further back the ancestors of hominins diverged from the ancestors of chimpanzees and bonobos (Langergraber et al., 2012). Whereas most phenotypic features that distinguish present-day humans from the apes and from their extinct hominin relatives are likely to be genetically complex, metabolic differences may sometimes have a comparatively simple genetic background because genomic changes that affect genes encoding enzymes may affect flux in metabolic pathways.

To find metabolic differences that set humans apart from their closest evolutionary relatives we investigated the metabolomes of the brain, muscle and kidney in humans, apes and monkeys. Whereas we find no such differences among the metabolites analyzed in muscle and kidney, in the brain, steady state concentrations of many compounds involved in amino acid metabolism are present in higher or lower levels in humans versus other primates. In the future, it may be of interest to investigate the consequences of these human-specific metabolic features for the synthesis and catabolism of amino acids.

Oxidative phosphorylation and purine biosynthesis stand out in that metabolites in both pathways are present in lower concentrations in humans than in the other primates analyzed. Whereas oxidative phosphorylation is lower in the three brain regions analysed but not in muscle and kidney, purine biosynthesis is decreased in all tissues analysed, although most drastically in brain.

Humans and apes diverged so long ago that almost every gene carries changes that potentially alter its function by affecting its regulation or the structure of the encoded gene product. In contrast, modern humans and Neandertals and Denisovans diverged so recently that for about 90% of the genome, the two archaic human groups fall within the variation of present-day humans (Green et al., 2010). Furthermore, when modern and archaic humans met about 50,000 years ago, they interbred. This resulted in that Neandertal DNA fragments that together make up about half of the Neandertal genome exist in present-day humans (Sankararaman et al., 2014). The number of proteins that carry amino acid substitutions in all or almost all humans that differ from Neandertals and apes is therefore only about one hundred (Pääbo, 2014). It is unclear if any of these substitutions have any functional consequences.

The alanine to valine substitution at position 429 in ADSL is one of this small number of substitutions. It is particularly likely to have functional consequence. Position 429 is conserved as alanine in most tetrapods suggesting that it may be of importance. Position 429 is also located only three positions away from position 426, where an arginine to histidine substitution causes the most common form of adenylosuccinase deficiency in present-day humans (Edery et al., 2003; Kmoch et al., 2000; Maaswinkel-Mooij et al., 1997; Marie et al., 1999; Race et al., 2000). Further evidence suggesting that a change in *ADSL* may have been of importance in the evolution of modern humans comes from a screen for genomic regions that have experienced selective sweeps in humans after their split from Neandertals but before the separation of Africans and Eurasians (Racimo, 2016). In that study, a genomic region centered around *ADSL* is among the top 20 candidate regions, although it contains also other genes. Furthermore, previous work has shown that the A429V substitution reduces the thermal stability of the ADSL protein *in vitro* (Van Laer et al., 2018). We therefore decided to analyze if it might be involved in the reduced purine biosynthesis seen in present-day humans by investigating the function of the ancestral, Neandertal-like and the derived, modern human-like forms of ADSL *in vitro* and *in vivo*.

We confirm the previous finding (Van Laer et al., 2018) that the A429V substitution does not affect the kinetic properties of the ADSL enzyme but decreases its thermal stability and show that the substitution also decreases the stability of the tetrameric complex of the enzyme. When introduced in the mouse ADSL protein in conjunction with a primate-specific substitution at the adjacent position 428, it reduces the enzymatic activity detected in nine tissues analyzed, most drastically in the brain, and results in a reduction in purine biosynthesis, thus recapitulating differences seen between humans and chimpanzees and macaques.

To investigate how the A429V substitution may affect the metabolism of human cells, we used CRISPR-*Cas9* to introduce the ancestral, Neandertal-like substitution into human cells. The concentrations of all nine metabolites detected downstream of ADSL in purine biosynthesis are increased in human cells carrying the ancestral substitution. In chimpanzee cells, the same nine metabolites are similarly increased. Notably, the expression of ADSL messenger RNA does not differ between human and chimpanzee cells, nor between wild type and ancestralized cells (not shown). Thus, the A429V substitution is responsible for much or all of the difference in purine biosynthesis observed when human tissues are compared to ape and monkey tissues, indicating that this change in metabolism occurred in humans after their separation from the ancestor shared with Neandertals and Denisovans.

An interesting question is what downstream effects a reduced purine biosynthesis in modern humans might have. In this regard, it is of interest that the A429V substitution correlates with reduction in phosphorylation of AMPK, a major regulator of cellular energy homeostasis. It is possible that reduced AMPK activity may contribute to the reduction in oxidative phosphorylation seen in human brains through its direct effects on this pathway (Lantier et al., 2014; Nam et al., 2016) and/or through its effects on the generation of mitochondria in cells (Bergeron et al., 2001; Jäger et al., 2007; Marin et al., 2017; Reznick et al., 2007). However, we find no evidence that oxidative phosphorylation is affected in the ancestralized human cells nor in the mice carrying the human substitution. Thus, if the A429V substitution contributes to a reduction in oxidative phosphorylation in humans, it presumably does so in conjunction with other genetic changes.

In general, it is interesting that although ADSL is expressed and functions in all tissues, the down-regulation of purine biosynthesis in humans relative to apes, and in humanized mice relative to wild-type mice, is most pronounced in the brain. It is also interesting that mutations in humans that affect enzymes involved in purine metabolism have more pathological consequences in the nervous system than in other organs (Fumagalli et al., 2017; Micheli et al., 2011). It is thus possible that the A429V substitution in ADSL has contributed to human-specific changes in brain development and function. Future work will have to address this and other possibilities.

## Acknowledgements

We are grateful to the NOMIS Foundation and the Max Planck Society for funding to SP; to the Bonfils-Stanton Foundation, the Butler Family Fund of the Denver Foundation, the Sam and Freda Davis Charitable Trust, and the Theodore T Puck Endowment of the University of Denver for funding to DP; to Stephane Peyregne and Karin Mörl for helpful input, to Rowina Voigtländer, Wulf Hevers and the animal facility staff for mouse breeding, and to Alexander Cagan and Wulf Hevers for tissue dissections. Animal breeding and experiments were done under the permission AZ: 24-9162.11/12/12 (T 10/14) from the Landesdirektion Sachsen.

## Methods

All methodological information, including the description of tissue sample sources and dissection, mouse genome editing, genome editing in human cells, enzyme essays, mass spectrometric metabolite measurements, protein properties assessment, and statistical analysis are described in detail in Supplementary Information.

## References

Ariyananda, L.D.Z., Lee, P., Antonopoulos, C., Colman, R.F., 2009. Biochemical and biophysical analysis of five disease-associated human adenylosuccinate lyase mutants. Biochemistry 48, 5291–5302. doi:10.1021/bi802321m

Bergeron, R., Ren, J.M., Cadman, K.S., Moore, I.K., Perret, P., Pypaert, M., Young, L.H., Semenkovich, C.F., Shulman, G.I., 2001. Chronic activation of AMP kinase results in NRF-1 activation and mitochondrial biogenesis. American Journal of Physiology - Endocrinology and Metabolism 281, E1340–6. doi:10.1152/ajpendo.2001.281.6.E1340

Brosius, J.L., Colman, R.F., 2002. Three subunits contribute amino acids to the active site of tetrameric adenylosuccinate lyase: Lys268 and Glu275 are required. Biochemistry 41, 2217–2226.

Castellano, S., Parra, G., Sánchez-Quinto, F.A., Racimo, F., Kuhlwilm, M., Kircher, M., Sawyer, S., Fu, Q., Heinze, A., Nickel, B., Dabney, J., Siebauer, M., White, L., Burbano, H.A., Renaud, G., Stenzel, U., Lalueza-Fox, C., la Rasilla de, M., Rosas, A., Rudan, P., Brajkovic, D., Kucan, Ž., Gušic, I., Shunkov, M.V., Derevianko, A.P., Viola, B., Meyer, M., Kelso, J., Andrés, A.M., Pääbo, S., 2014. Patterns of coding variation in the complete exomes of three Neandertals. Proc Natl Acad Sci USA 111, 6666–6671. doi:10.1073/pnas.1405138111

Ciardo, F., Salerno, C., Curatolo, P., 2001. Neurologic aspects of adenylosuccinate lyase deficiency. J. Child Neurol. 16, 301–308. doi:10.1177/088307380101600501

Daignan-Fornier, B., Pinson, B., 2012. 5-Aminoimidazole-4-carboxamide-1-beta-D-ribofuranosyl 5’-Monophosphate (AICAR), a Highly Conserved Purine Intermediate with Multiple Effects. Metabolites 2, 292–302. doi:10.3390/metabo2020292

Davies, S.P., Helps, N.R., Cohen, P.T., Hardie, D.G., 1995. 5’-AMP inhibits dephosphorylation, as well as promoting phosphorylation, of the AMP-activated protein kinase. Studies using bacterially expressed human protein phosphatase-2C alpha and native bovine protein phosphatase-2AC. FEBS Lett 377, 421–425. doi:10.1016/0014-5793(95)01368-7

Edery, P., Chabrier, S., Ceballos-Picot, I., Marie, S., Vincent, M.-F., Tardieu, M., 2003. Intrafamilial variability in the phenotypic expression of adenylosuccinate lyase deficiency: a report on three patients. Am J Med Genet A 120A, 185–190. doi:10.1002/ajmg.a.20176

Fumagalli, M., Lecca, D., Abbracchio, M.P., Ceruti, S., 2017. Pathophysiological Role of Purines and Pyrimidines in Neurodevelopment: Unveiling New Pharmacological Approaches to Congenital Brain Diseases. Front. Pharmacol. 8, 941. doi:10.3389/fphar.2017.00941

Giavalisco P., Li Y., Matthes A., Eckhardt A., Hubberten H. M., Hesse H., Segu S., Hummel J., Köhl K., Willmitzer L. (2011). Elemental formula annotation of polar and lipophilic metabolites using (13) C, (15) N and (34) S isotope labelling, in combination with high-resolution mass spectrometry. Plant J. 10.1111/j.1365-313X.2011.04682.x

Green, R.E., Krause, J., Briggs, A.W., Maricic, T., Stenzel, U., Kircher, M., Patterson, N., Li, H., Zhai, W., Fritz, M.H.-Y., Hansen, N.F., Durand, E.Y., Malaspinas, A.-S., Jensen, J.D., Marques-Bonet, T., Alkan, C., Prüfer, K., Meyer, M., Burbano, H.A., Good, J.M., Schultz, R., Aximu-Petri, A., Butthof, A., Höber, B., Höffner, B., Siegemund, M., Weihmann, A., Nusbaum, C., Lander, E.S., Russ, C., Novod, N., Affourtit, J., Egholm, M., Verna, C., Rudan, P., Brajkovic, D., Kucan, Ž., Gušic, I., Doronichev, V.B., Golovanova, L.V., Lalueza-Fox, C., la Rasilla de, M., Fortea, J., Rosas, A., Schmitz, R.W., Johnson, P.L.F., Eichler, E.E., Falush, D., Birney, E., Mullikin, J.C., Slatkin, M., Nielsen, R., Kelso, J., Lachmann, M., Reich, D., Pääbo, S., 2010. A Draft Sequence of the Neandertal Genome. Science 328, 710–722. doi:10.1126/science.1188021

Hawley, S.A., Selbert, M.A., Goldstein, E.G., Edelman, A.M., Carling, D., Hardie, D.G., 1995. 5’-AMP activates the AMP-activated protein kinase cascade, and Ca2+/calmodulin activates the calmodulin-dependent protein kinase I cascade, via three independent mechanisms. J Biol Chem 270, 27186–27191. doi:10.1074/jbc.270.45.27186

Jaeken, J., Van Den Berghe, G., 1984. An infantile autistic syndrome characterised by the presence of succinylpurines in body fluids. Lancet 2, 1058–1061.

Jäger, S., Handschin, C., Pierre, J.S., Spiegelman, B.M., 2007. AMP-activated protein kinase (AMPK) action in skeletal muscle via direct phosphorylation of PGC-1α. Proc Natl Acad Sci USA 104, 12017–12022. doi:10.1073/pnas.0705070104

Jurecka, A., Jurkiewicz, E., Tylki-Szymańska, A., 2012. Magnetic resonance imaging of the brain in adenylosuccinate lyase deficiency: a report of seven cases and a review of the literature. Eur. J. Pediatr. 171, 131–138. doi:10.1007/s00431-011-1503-9

Jurecka, A., Zikánová, M., Kmoch, S., Tylki-Szymańska, A., 2015. Adenylosuccinate lyase deficiency. J Inherit Metab Dis 38, 231–242. doi:10.1007/s10545-014-9755-y

Kim, J., Yang, G., Kim, Y., Kim, J., Ha, J., 2016. AMPK activators: mechanisms of action and physiological activities. Exp. Mol. Med. 48, e224–e224. doi:10.1038/emm.2016.16

Kmoch, S., Hartmannova, H., Stiburkova, B., Krijt, J., Zikanova, M., Sebesta, I., 2000. Human adenylosuccinate lyase (ADSL), cloning and characterization of full-length cDNA and its isoform, gene structure and molecular basis for ADSL deficiency in six patients. Hum Mol Genet 9, 1501–1513.

Langergraber, K.E., Prüfer, K., Rowney, C., Boesch, C., Crockford, C., Fawcett, K., Inoue, E., Inoue-Muruyama, M., Mitani, J.C., Muller, M.N., Robbins, M.M., Schubert, G., Stoinski, T.S., Viola, B., Watts, D., Wittig, R.M., Wrangham, R.W., Zuberbühler, K., Pääbo, S., Vigilant, L., 2012. Generation times in wild chimpanzees and gorillas suggest earlier divergence times in great ape and human evolution. Proceedings of the National Academy of Sciences 109, 15716–15721. doi:10.1073/pnas.1211740109

Lantier, L., Fentz, J., Mounier, R., Leclerc, J., Treebak, J.T., Pehmøller, C., Sanz, N., Sakakibara, I., Saint-Amand, E., Rimbaud, S., Maire, P., Marette, A., Ventura-Clapier, R., Ferry, A., Wojtaszewski, J.F.P., Foretz, M., Viollet, B., 2014. AMPK controls exercise endurance, mitochondrial oxidative capacity, and skeletal muscle integrity. The FASEB Journal 28, 3211–3224. doi:10.1096/fj.14-250449

Lee, P., Colman, R.F., 2007. Expression, purification, and characterization of stable, recombinant human adenylosuccinate lyase. 51, 227–234. doi:10.1016/j.pep.2006.07.023

Lisec J, Schauer N, Kopka J, Willmitzer L,Fernie AR (2006) Gas chromatography mass spectrometry-based metabolite profiling in plants. Nat Protoc 1:387–396

Maaswinkel-Mooij, P.D., Laan, L.A., Onkenhout, W., Brouwer, O.F., Jaeken, J., Poorthuis, B.J., 1997. Adenylosuccinase deficiency presenting with epilepsy in early infancy. J Inherit Metab Dis 20, 606–607. doi:10.1023/A:1005323512982

Marie, S., Cuppens, H., Heuterspreute, M., Jaspers, M., Tola, E.Z., Gu, X.X., Legius, E., Vincent, M.F., Jaeken, J., Cassiman, J.J., Van Den Berghe, G., 1999. Mutation analysis in adenylosuccinate lyase deficiency: eight novel mutations in the re-evaluated full ADSL coding sequence. Hum. Mutat. 13, 197–202. doi:10.1002/(SICI)1098-1004(1999)13:3<197::AID-HUMU3>3.0.CO;2-D

Marie, S., Heron, B., Bitoun, P., Timmerman, T., Van Den Berghe, G., Vincent, M.-F., 2004. AICA-ribosiduria: a novel, neurologically devastating inborn error of purine biosynthesis caused by mutation of ATIC. Am J Hum Genet 74, 1276–1281. doi:10.1086/421475

Marin, T.L., Gongol, B., Zhang, F., Martin, M., Johnson, D.A., Xiao, H., Wang, Y., Subramaniam, S., Chien, S., Shyy, J.Y.J., 2017. AMPK promotes mitochondrial biogenesis and function by phosphorylating the epigenetic factors DNMT1, RBBP7, and HAT1. Sci Signal 10. doi:10.1126/scisignal.aaf7478

Micheli, V., Camici, M., Tozzi, M.G., Ipata, P.L., Sestini, S., Bertelli, M., Pompucci, G., 2011. Neurological disorders of purine and pyrimidine metabolis. Curr Top Med Chem 11, 923–947.

Mihaylova, M.M., Shaw, R.J., 2011. The AMPK signalling pathway coordinates cell growth, autophagy and metabolism. Nature Cell Biology 13, 1016–1023. doi:10.1038/ncb2329

Nam, K., Oh, S., Shin, Incheol, 2016. Ablation of CD44 induces glycolysis-to-oxidative phosphorylation transition via modulation of the c-Src–Akt–LKB1–AMPKα pathway. Biochem J 473, 3013.

Pääbo, S., 2014. The human condition-a molecular approach. Cell 157, 216–226. doi:10.1016/j.cell.2013.12.036

Phillips, P.C., 2008. Epistasis--the essential role of gene interactions in the structure and evolution of genetic systems. Nat. Rev. Genet. 9, 855–867. doi:10.1038/nrg2452

Prüfer, K., Racimo, F., Patterson, N., Jay, F., Sankararaman, S., Sawyer, S., Heinze, A., Renaud, G., Sudmant, P.H., de Filippo, C., Li, H., Mallick, S., Dannemann, M., Fu, Q., Kircher, M., Kuhlwilm, M., Lachmann, M., Meyer, M., Ongyerth, M., Siebauer, M., Theunert, C., Tandon, A., Moorjani, P., Pickrell, J., Mullikin, J.C., Vohr, S.H., Green, R.E., Hellmann, I., Johnson, P.L.F., Blanche, H., Cann, H., Kitzman, J.O., Shendure, J., Eichler, E.E., Lein, E.S., Bakken, T.E., Golovanova, L.V., Doronichev, V.B., Shunkov, M.V., Derevianko, A.P., Viola, B., Slatkin, M., Reich, D., Kelso, J., Pääbo, S., 2014. The complete genome sequence of a Neanderthal from the Altai Mountains. Nature 505, 43–49. doi:10.1038/nature12886

Race, V., Marie, S., Vincent, M.F., Van Den Berghe, G., 2000. Clinical, biochemical and molecular genetic correlations in adenylosuccinate lyase deficiency. Hum Mol Genet 9, 2159–2165. doi:10.1093/hmg/9.14.2159

Racimo, F., 2016. Testing for ancient selection using cross-population allele. Genetics 202, 733–750. doi:10.1534/genetics.115.178095/-/DC1

Ray, S.P., Deaton, M.K., Capodagli, G.C., Calkins, L.A.F., Sawle, L., Ghosh, K., Patterson, D., Pegan, S.D., 2012. Structural and biochemical characterization of human adenylosuccinate lyase (ADSL) and the R303C ADSL deficiency-associated mutation. Biochemistry 51, 6701–6713. doi:10.1021/bi300796y

Reznick, R.M., Zong, H., Li, J., Morino, K., Moore, I.K., Yu, H.J., Liu, Z.-X., Dong, J., Mustard, K.J., Hawley, S.A., Befroy, D., Pypaert, M., Hardie, D.G., Young, L.H., Shulman, G.I., 2007. Aging-associated reductions in AMP-activated protein kinase activity and mitochondrial biogenesis. Cell Metabolism 5, 151–156. doi:10.1016/j.cmet.2007.01.008

Riesenberg, S., Chintalapati, M., Macak, D., Kanis, P.,Maricic, T., Pääbo, S., 2019. Simultaneous precise editing of multiple genes in human cells, Nucleic Acids Research, Volume 47, Issue 19, 04 November 2019, Page e116, https://doi.org/10.1093/nar/gkz669

Sankararaman, S., Mallick, S., Dannemann, M., Prüfer, K., Kelso, J., Pääbo, S., Patterson, N., Reich, D., 2014. The genomic landscape of Neanderthal ancestry in present-day humans. Nature 507, 354–357. doi:10.1038/nature12961

Spiegel, E.K., Colman, R.F., Patterson, D., 2006. Adenylosuccinate lyase deficiency. Mol Genet Metab 89, 19–31. doi:10.1016/j.ymgme.2006.04.018

Sugimoto, M., Sakagami, H., Yokote, Y., Onuma, H., Kaneko, M., Mori, M., Sakaguchi, Y., Soga, T., Tomita, M., 2012. Non-targeted metabolite profiling in activated macrophage secretion. Metabolomics 8, 624–633. doi:10.1007/s11306-011-0353-9

Tomasello, M., 2019. Becoming Human: A Theory of Ontogeny. Harvard University Press.

Van Laer, B., Kapp, U., Soler-Lopez, M., Moczulska, K., Pääbo, S., Leonard, G., Mueller-Dieckmann, C., 2018. Molecular comparison of Neanderthal and Modern Human adenylosuccinate lyase. Sci. Rep. 8, 18008. doi:10.1038/s41598-018-36195-5

Varki, A., Altheide, T.K., 2005. Comparing the human and chimpanzee genomes: searching for needles in a haystack. Genome Res 15, 1746–1758. doi:10.1101/gr.3737405

Vliet, L.K., Wilkinson, T.G., Duval, N., Vacano, G., Graham, C., Zikánová, M., Skopova, V., Baresova, V., Hnízda, A., Kmoch, S., Patterson, D., 2011. Molecular characterization of the AdeI mutant of Chinese hamster ovary cells: a cellular model of adenylosuccinate lyase deficiency. Mol Genet Metab 102, 61–68. doi:10.1016/j.ymgme.2010.08.022

Zikánová, M., Skopova, V., Hnízda, A., Krijt, J., Kmoch, S., 2010. Biochemical and structural analysis of 14 mutant adsl enzyme complexes and correlation to phenotypic heterogeneity of adenylosuccinate lyase deficiency. Hum. Mutat. 31, 445–455. doi:10.1002/humu.21212

