## Supplemental Info for "Reduced purine biosynthesis in humans after their divergence from Neandertals"

**Supplementary Information**

**Supplementary Methods**

**Samples.** Human samples were obtained from the Netherlands Brain Bank. Amsterdam. Netherlands. Written consent for the use of human tissues for research was obtained from all donors or their next of kin. All subjects were defined as healthy controls by forensic pathologists at the brain bank. All subjects suffered sudden death with no prolonged agonal state. Chimpanzee samples were from the tissue collection of the Max Planck Institute for Anthropology. Rhesus macaque samples were obtained from the Suzhou Experimental Animal Center. China. All chimpanzees and rhesus macaques used in this study suffered sudden deaths for reasons other than their participation in this study and without any relation to the tissue used. Prefrontal cortex (PFC) dissections were made from the frontal part of the superior frontal gyrus. For all samples. we took special care to dissect gray matter only.

**Genome editing in mice.** Male pronuclei of C57BL/6NCrl fertilized oocytes were injected with a mix consisting of Cas9 or Cas9-D10A Nickase mRNA (50 ng/µl). gRNA in form of RNA (20 ng/µl) and single strand oligonucleotide (50 ng/µl). The sequence of the 99mer oligonucleotide is 5’-GTGGTCAAGCAGGAAGGAGGTGACAATGACCTTATAGAGCGCATC**CAGGTT**GACGCCTACTTCAGCCCCATCCACTCACAGCTGGAGCACTTGCTGGAC-3’. The nucleotide substitutions for changing the amino acids arginine (CGG) and alanine (GCA) to glutamine (CAG) and valine (GTT) that occur in the human sequence are depicted in bold. The humanized sequence introduces two new restriction sites for *BstNI* and *HincII* that facilitate genotyping of the founders. The sequence of the two guide RNAs are: Adsl-gRNA10: 5’-GGCTGAAGTAGGCATCTGCC-3’ for Cas9 and Adsl-gRNA15: 5’-CTCACAGCTGGAGCACTTGC-3’ used together with gRNA10 for the Cas9-D10A injections. After injections. the surviving embryos were transferred into Crl:CD1(ICR) pseudopregnant recipient female mice (20–25 embryos per recipient). Tail DNA of the founders was PCR amplified with Adsl-F: 5’-CGATGTCTGACTGTAAGGTCTAC-3’ and Adsl-R: 5’-CCAGAGGCTCAGGCTCTGCATCA-3’ and digested with *HincII*. In total 6 from 105 born mice were positive for the humanized sequence.

**Genome editing in human cells.** 409-B2 human induced pluripotent stem cells (hiPSCs) with a doxycycline inducible Cas9 nickase (D10A mutation) and enhanced homology-directed repair efficiency (DNA-PKcs K3753R mutation) were incubated with mTesR1 medium (StemCell Technologies. 05851) containing 2 µg/ml doxycycline (Clontech. 631311) two days prior to lipofection of a gRNA (duplex of chemically synthesized crRNA and tracrRNA. alt-CRISPR IDT) pair and single stranded DNA donor as described (Riesenberg et al.. 2019). Lipofection by RNAiMAX (Invitrogen. 13778075) was done using a final concentration of 7.5 nM of each gRNA (ADSL_target1: CAACCTGGATACGCTCTATG. ADSL_target2: CAGTCCCATTCACTCCCAGT) and 10 nM of single stranded DNA donor (ADSL_V429A: CAGCAGGCAGCTTCTGTGGTTAAGCAGGAAGGGGGTGACAATGACCTCAT*T*GAGCGTATCCAGG**C**TGATGCCTACTTCAGTCCCATTCACTC*T*CAGTTGGATCATTTACTGGATCCTTCTTCTTTCACTGGTCGTGCCTCCCAG; ancestral mutation = bold. silent mutations to prevent recutting: italic). The lipofection mix was exchanged to regular mTesR1 medium after 24h and cells were propagated. After dissociation using Accutase (SIGMA. A6964) cells were plated in a single cell dilution that gave rise to single cell derived colonies. DNA isolation. Illumina library preparation and sequencing analysis to conform editing success was done as described (Riesenberg et al.. 2019). Primers for DNA target amplification were ADSL_forward: AAAGTGTTCAAACGCCCTGT and ADSL_reverse: AGAGAGTACCCCAGGTTCTGC.

**Enzyme Assays.** Tissue samples were homogenized in 300 mM sucrose. 10 mM Tris pH 7.4. 10 mM EDTA using a PTFE tissue grinder and mixer motor. Samples were centrifuged at 4 °C for 30 minutes in a microfuge at maximum speed. Supernatant was collected and stored at -70 °C. Protein concentration was determined using a bicinchoninic acid assay kit (Sigma) and a BioTek PowerWave XS2 plate reader. measuring absorbance at λ 562 nm. Absorbance values were fitted against a BSA standard curve (0. 12.5. 25. 50. 75. 100. 150. 200 μg/ml). For the ADSL assays. 50 ng protein was added into a 60 μM adenylosuccinate (AMPS)/40 mM Tris (pH 7.4)/10% glycerol solution. and conversion of adenylosuccinate to AMP was monitored by UV spectrophotometry. measuring absorption at λ 282 nm every minute for 20 minutes (Greiner UV-Star® 96 well plates and BioTek PowerWave XS2 plate reader). ADSL activity was calculated using the extinction coefficient for the reaction (10.000 M-1 cm-1) in Excel. Wilcoxon tests and box plots were performed using R.

**CE-MS measurements of metabolomes.** For each sample. metabolites were extracted from the frozen tissue powder by 1 ml methanol containing 20 µM each of L-Methionine sulfone. 2-morpholinoethanesulfonic acid. monohydrate. and sodium d-camphor-10-sulfonic acid. 500 µl of the lysate was transferred to an Eppendorf tube containing 500 µl chloroform and 200 µl of Milli-Q water. After 30 seconds of vortexing and 15 min of centrifugation at 4 °C. 300 µl of the aqueous phase was concentrated and dried to complete dryness in an ultrafiltration tube (Millipore) followed by speed vacuum for 3 hours at 35 °C. The dried samples were mixed with 100 µl of Milli-Q water containing 100 µM each of 3-aminopyrrolidine and trimesate. and filtered with an ultrafiltration tube (Millipore) at 9.100 × g for 2 hours at 4 °C immediately before 7 µl was used in a CE-TOF-MS (Agilent Technologies) to detect cationic metabolites and anionic metabolites. The instrumentation and measurement conditions used for CE-TOF-MS were according to (Sugimoto et al.. 2012).

The in-house software MasterHands was used for peak detection. time alignment. and peak area integration. The intensities of each metabolite in the samples were calculated based on the comparison of peak area normalized by internal standards added to the sample as well as external standards. Metabolites detected in less than half of the samples were excluded from the following analysis.

**GC-MS measurements of metabolomes.** Metabolites were extracted from the frozen tissue powder by a methanol:water:chloroform (2.5:1:1 (v/v/v)) extraction. In brief. 100mg of frozen powdered tissue material was resuspended 1 mL extraction solution containing 0.1 µg mL-1 of U-13C6-sorbitol. The samples were incubated for 10 min at 4 °C on an orbital shaker. This step was followed by ultrasonication in a bath-type sonicator for 10 min at room temperature. Finally. the insoluble tissue material was pelleted by a centrifugation step (5 min; 14.000g). and the supernatant transferred to a fresh 2 mL Eppendorf tube. To separate the organic from the aqueous phase. 300 µL H2O and 300 µL chloroform were added to the supernatant. vortexed. and centrifuged (2 min; 14.000g). Subsequently. 200 µL of the upper. aqueous phase was collected and concentrated to complete dryness in a speed vacuum at room temperature. Extract derivatization and GC-MS measurements were performed according to Lisec J (2006).

MS peaks were aligned across samples and annotated to known compounds according to Giavalisco et al.. 2011. Unannotated MS peaks were excluded. and each tissue was further analyzed separately. The obtained metabolite concentration values (apex height of the quantitative compound identifier mass) were normalized within each sample to the abundance of an internal standard (13C sorbitol). and log10 transformed. To avoid negative values. metabolite/standard ratios were scaled up by factor 3.000 prior to log10 transformation. Next. we filtered out metabolites containing more or equal to 50% missing values. To normalize distributions of values between samples. we additionally divided intensity values for the remaining metabolites to the upper quartiles.

**LC-MS sample preparation.** Adhered cells were grown mTesR1 medium (StemCell Technologies. 05851) in 6-well plates. After counting. medium solution was gently discarded and cells were quickly washed twice with 1 ml of 1x PBS. Cells were harvested in PBS. placed into 2 ml Eppendorf (safe lock) tubes. suspended by slight pipetting. pelleted by centrifugation at 4 °C and snap-frozen in liquid nitrogen. Pellets were stored at -80 °C. The several blank samples were added to the end of each batch (48 samples). They were 2 ml Eppendorf (safe lock) tubes without any sample material but subjected to all extraction procedures. Cells were quickly resuspended in 75 μl of water on ice and metabolites were extracted by the after addition of 1 ml of -20 °C cold methanol: methyl-tert-butyl-ether (1∶3 [v/v]) extraction buffer. vortexed. and sonicated for 10 min in an ice-cooled sonication bath followed by incubation for a 30 min at 4 °C on an orbital shaker as described (Giavalisco et al. 2011). 700 μl of a H2O:methanol mix (3:1 (v/v)). containing 5.0 μg of GMP-15N5 and Methionine-methyl-13C.d3 was added and the mixture was incubated for a 10 min at 4 °C on an orbital shaker. To separate the organic and the aqueous phases and precipitate proteins the mixture was spun down for 10 min at 15.000 x g at 4 °C. Subsequently. 300 μl of the lower aqueous phase containing hydrophilic compounds was transferred to a 1.5 ml Eppendorf tube and concentrated to complete dryness in a speed vacuum centrifuge at room temperature. Dry metabolic extracts were stored at -80 °C prior to mass spectrometry analysis.

In the LC-MS datasets. in addition to the individual cells samples. we measured mixtures of samples (pooled extracted samples. TQC) after every 10th sample. providing information on system performance (sensitivity and retention time consistency). sample reproducibility. and compound stability over the time of the MS-based analysis.

**Metabolite abundance measurements.** The dried extracts were resuspended in 100 µl of ice-cold 20% aqueous solution of acetonitrile prior to mass spectrometry analysis. After brief rigorous vortexing. the samples were incubated for 30 min at 4 °C on an orbital shaker followed by a 10 min ultra-sonication in an ice-cooled sonication bath and centrifugation for 10 min at 15.000 x g at 4 °C. For mass spectrometry analysis. 40 µl of supernatant was transferred to a 350 μl autosampler glass vials (Glastechnik Grafenroda. Germany). Chromatographic separation of metabolites prior to mass spectrometry was performed using Acquity I-Class UPLC system (Waters. UK). Metabolites were separated on a normal phase unbounded silica column RX-SIL (100 mm x 2.1 mm. 1.8 µm. Agilent. US) coupled to a guard precolumn with the same phase parameters. The mobile phases used for the chromatographic separation were water containing 10 mM ammonium acetate. 0.2 mM ammonium hydroxide in water:acetonitrile (95: 5(v:v)) mixture (buffer A) (pH value 8.0) and 100% acetonitrile (buffer B). The gradient separation was: 0 min 0% buffer A. 0.01-15 min linear gradient from 0% to 100% buffer A. 15-18 min 100% buffer A. 18-19 min linear gradient from 100% buffer A to 0% buffer A. and 19-32 min 0% buffer A. After 1 min washing with 100% buffer A the column was re-equilibrated with 100% buffer B. The flow rate was set to 500 µl/min. The column temperature was maintained at 32 °C. The mass spectra were acquired in positive and negative mode using a heated electrospray ionization source in combination with Q Exactive Hybrid Quadrupole-Orbitrap mass spectrometer (Thermo Scientific. Germany). Negative ion mode samples were run after the positive ion mode cohort with 6 µl injection of non-diluted samples. MS settings in positive acquisition mode: spray voltage was set to 4.5 kV in positive mode and to 3 kV in negative acquisition mode. S-lens RF level at 70. and heated capillary at 250°C; aux gas heater temperature was set at 350 °C; sheath gas flow rate was set to 45 arbitrary units; aux gas flow rate was set to 10 arbitrary units; sweep gas flow rate was set to 4 arbitrary units. Full scan resolutions were set to 70 000 at m/z 200. Full scan target was 10e6 with a maximum fill time of 50 ms. The spectra were recorded using full scan mode. covering a mass range from 100– 1500 m/z.

For quality control (TQC). a pooled sample of all metabolic extracts was prepared and injected 4 times before initiating the runs in order to condition the column. at least 4 times after each sub-cohort. and after the completion of the runs. In addition. the TQC sample was injected every 48 sample injections to assess instrument stability and reproducibility.

**Statistical analysis of metabolomes.** We performed principal component analysis (PCA) for each tissue separately in order to identify outlier samples. For the PCA analysis. the “prcomp” function in the R package “stats” was used. To identify metabolites with significant intensity changes. we implemented the Student’s t-test. A permutation procedure was used to assess the false positive rate. Briefly. we shuffled sample labels. applied t-test to two random groups of samples. and repeated this procedure 1.000 times. Then. a permutation p-value was used to estimate the ratio of permutations resulting in an equal or greater number of metabolites with significant intensity changes at a nominal significance cutoff of 0.05. To determine the probability of increase or decrease of metabolite intensities. we performed a binomial test. Log10 fold change of metabolite intensities was calculated as the average logarithm of intensity differences between two sample groups.

**Protein purification and dual substrate specific activity measurement.** ADSL protein versions were purified as described previously (Ray et al.. 2012). The specific activity in presence of single and double substrates shown in Figure 6 was measured using a customized HPLC system as described previously (Ray et al.. 2013).

**Native Gel Electrophoresis.** Aliquots of ADSL protein were thawed and equilibrated for 2-4h at room temperature. Subsequently. they were mixed with guanidine hydrochloride to final concentrations of 1.5M. 1.125M. 1.0125M. 0.9M. and 0.75MSamples were incubated overnight at room temperature and analyzed using native gel electrophoresis (Invitrogen. BN2111BX10. NativePage 4-16%) according to manufacturer’s manual. ImageJ was used for densitometry analysis of monomeric (M) and tetrameric (T) band of gel pictures where person assigning the peaks was blinded for the sample identity. Tetramer fraction was subsequently calculated using the following formula: T/(M+T). Differences between human and Neandertal-like versions of ADSL were calculated for each concentration of guanidine hydrochloride over 5 repetitions. Bonferroni correction was used to correct for number of comparisons (5 different guanidine hydrochloride concentrations).

**Circular Dichroism measurements.** ADSL versions were diluted in storage buffer as described to concentration of 0.2 mg/ml. Samples were thawed and equilibrated at room temperature for a minimum of 30min prior to the scan. Each sample was transferred to a 1mm cuvette for analysis. To analyse secondary structure. each sample was first scanned for wavelengths 200-250 nm and subsequently monitored at 222 nm from 50 to 80 °C using a Jasco J-815 Spectropolarimeter. Each ADSL protein version was analysed using a minimum of two separate protein batches and a minimum of two scans to avoid batch effects. Denaturation midpoint was designated as the temperature value closest to the 50% folded protein mark for each run.

**Supplementary Figures**

**Supplementary Figure 1.**

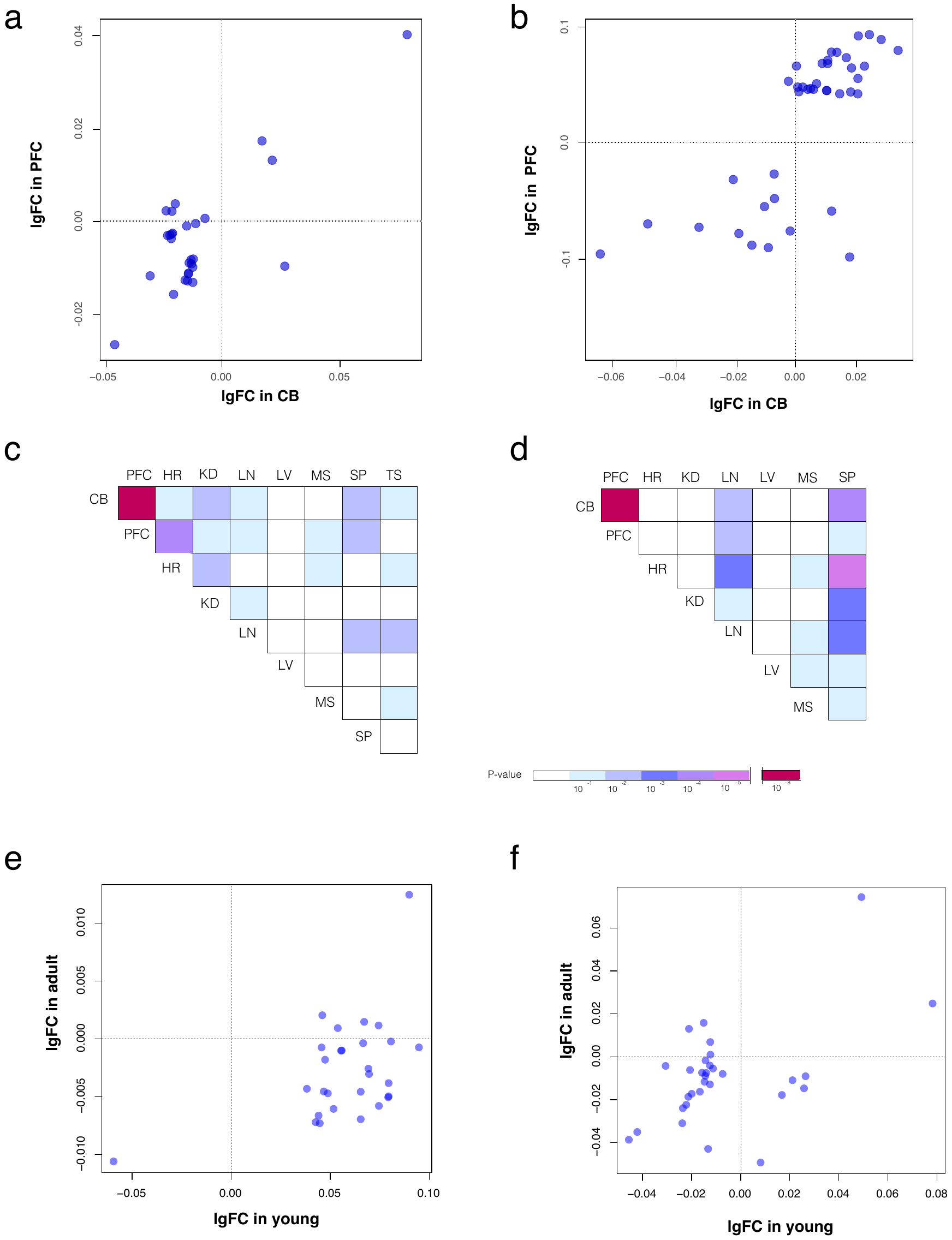

**Supplementary Figure 1. Correlations of metabolite concentration differences in mouse tissues. a.** Correlation of metabolite differences (given as log fold change) between wild type mice and mice carrying the V429A and R428Q substitutions in the ADSL protein in 12 week-old mice. **b.** The same correlations in 1 week-old mice. **c.d.** Correlations of metabolite concentration differences between cerebral cortex and other tissues in 12 week-old (c) and 1 week-old (d) mice. **e.** Correlation of metabolite concentration differences between 1 week- and 12 week-old mice in cerebral cortex. **f.** Correlation of in metabolite concentration differences between 1 week-old and 12 week-old mice in cerebellum.

**Supplementary Figure 2.**

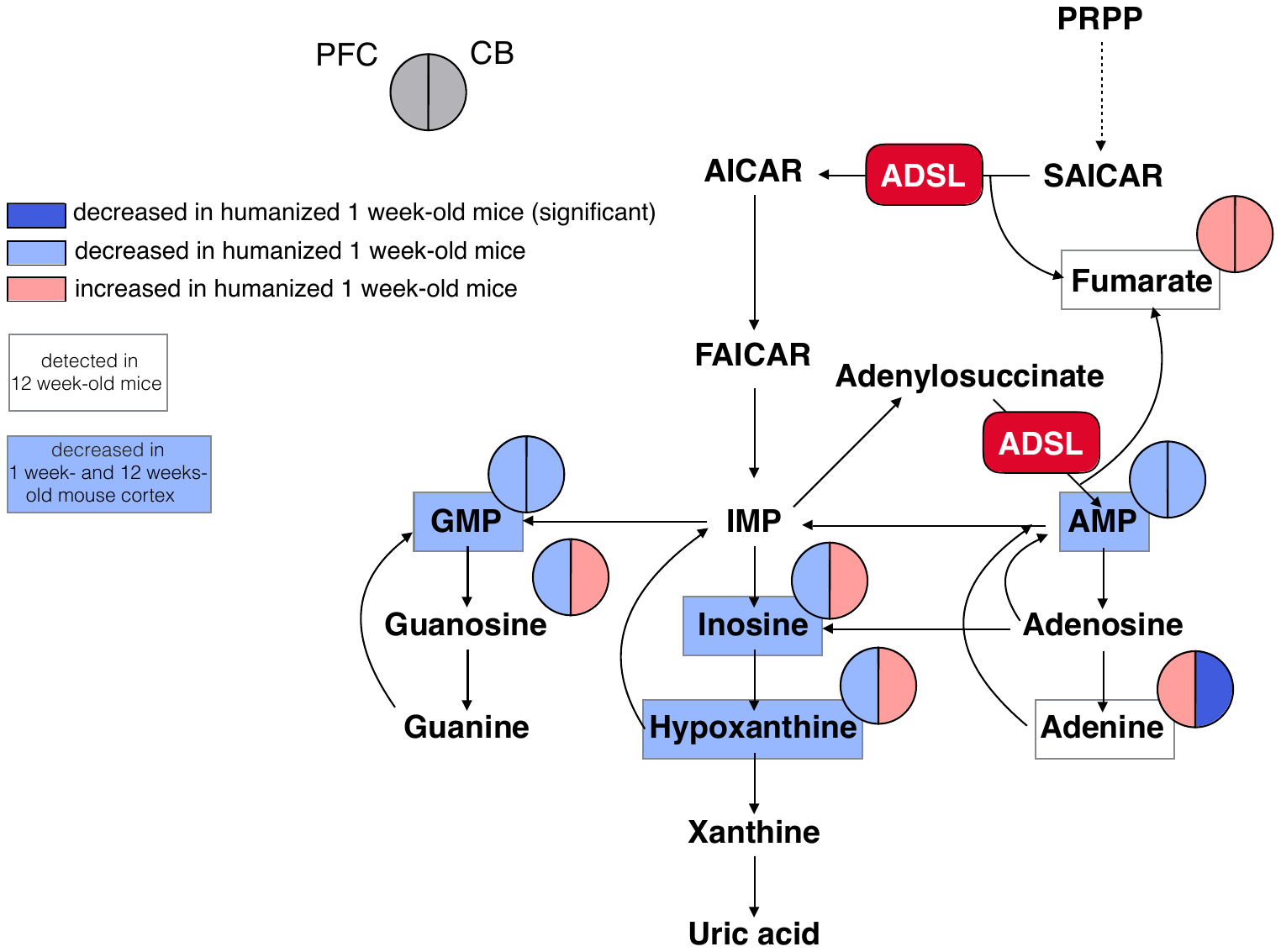

**Supplementary Figure 2.** **Purine biosynthesis in mice “humanized” pups for ADSL.** Pathway sketch showing changes in metabolite concentrations in 1 week-old mice carrying the V429A and R428Q substitutions in the ADSL protein. as well as the six metabolites detected and the four of these that are decreased in the cortex also 12 week-old humanized mice. PFC – prefrontal cortex. CB – cerebellum.

**Supplementary Figure 3.**
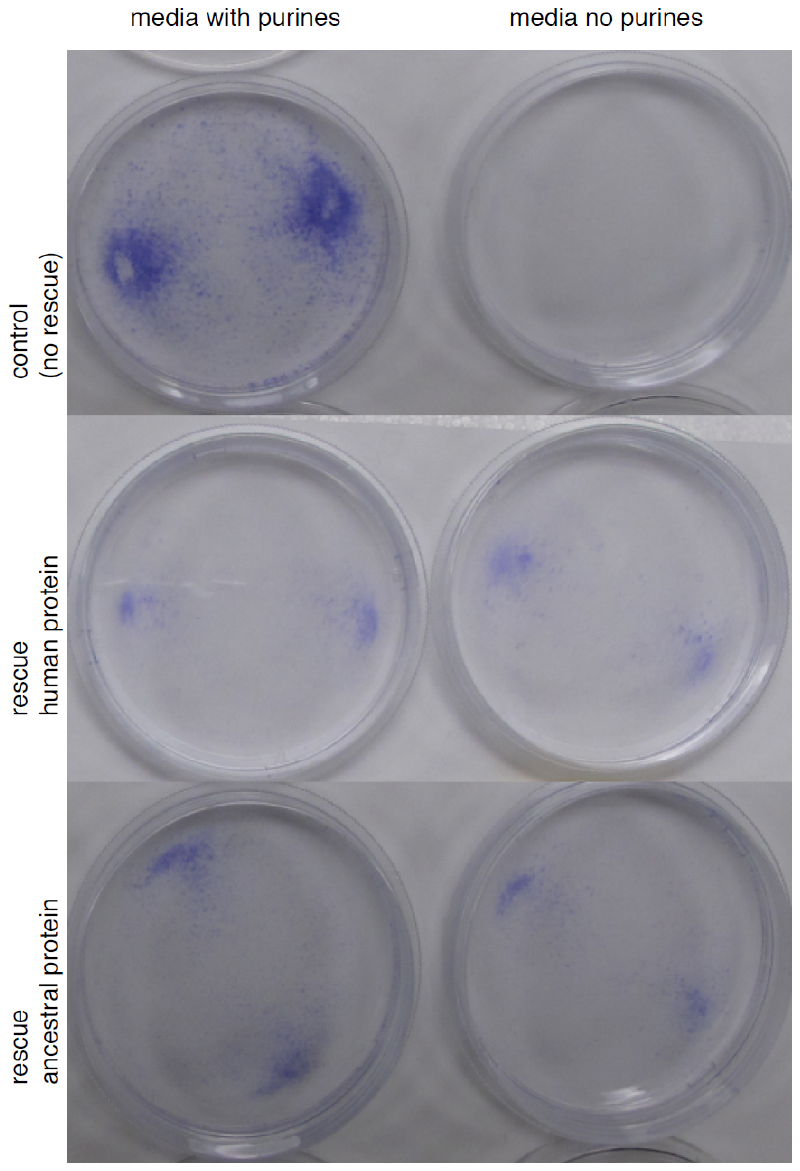

**a**

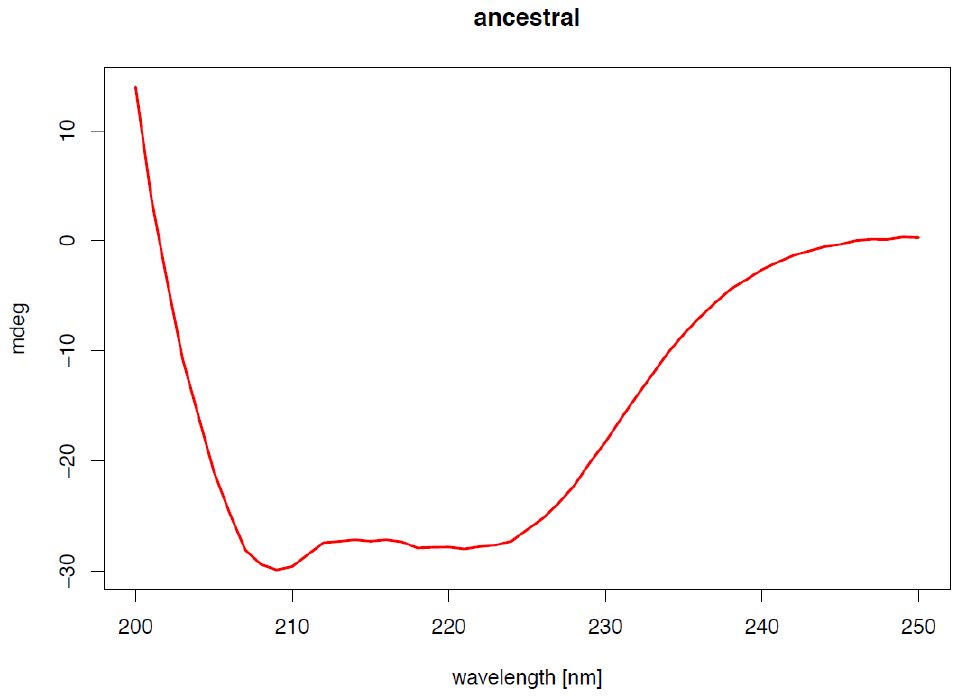

**b**

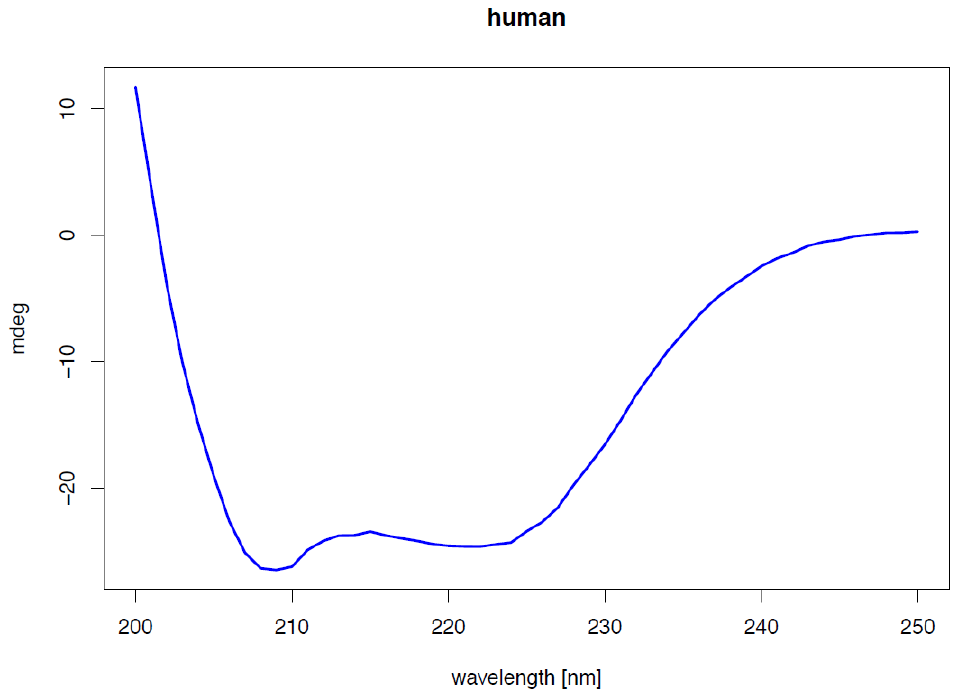

**Supplementary Figure 3. a.** Rescue of *Ade* cells in purine-free medium by Neandertal-like and modern human ADSL. **b.** CD spectra of ancestral. Neandertal-like and modern human ADSL.

**Supplementary Table 2. Metabolites showing concentration difference in humans.**

**a**

|  | **pval.adj.chimp** | **pval.adj.mac** | **FC.chimp** | **FC.mac** |
| --- | --- | --- | --- | --- |
| Gly | 3.14E-03 | 2.82E-03 | 3.8 | 4.2 |
| Ala | 1.82E-03 | 2.82E-03 | 3.0 | 3.4 |
| Ser | 1.45E-02 | 1.32E-02 | 4.6 | 3.4 |
| Pro | 6.84E-03 | 7.22E-03 | 8.8 | 11.4 |
| Val | 2.16E-03 | 3.96E-03 | 6.6 | 13.8 |
| Thr | 5.08E-03 | 7.22E-03 | 8.4 | 8.1 |
| Ile | 4.75E-03 | 6.38E-03 | 7.3 | 16.1 |
| Leu | 2.05E-03 | 3.64E-03 | 7.7 | 17.7 |
| Asn | 4.75E-03 | 2.82E-03 | 2.8 | 4.0 |
| Ornithine | 2.24E-02 | 1.53E-02 | 5.1 | 15.3 |
| Lys | 3.14E-03 | 3.73E-03 | 3.5 | 7.7 |
| Met | 7.28E-03 | 7.22E-03 | 8.5 | 12.8 |
| His | 1.30E-02 | 7.22E-03 | 4.1 | 6.4 |
| Methionine sulfoxide | 2.16E-03 | 3.64E-03 | 11.8 | 6.5 |
| Arg | 6.84E-03 | 7.22E-03 | 5.1 | 12.1 |
| Tyr | 6.84E-03 | 4.46E-03 | 4.2 | 8.8 |
| Trp | 2.05E-03 | 2.82E-03 | 3.2 | 6.4 |
| Cysteine-glutathione disulphide-divalent | 4.14E-02 | 2.99E-02 | 6.5 | 28.0 |
| 1-Methyladenosine | 1.30E-02 | 8.29E-03 | 6.3 | 16.2 |
| 5-Oxoproline | 3.80E-02 | 4.93E-02 | 3.2 | 2.4 |
| N-Acetylglucosamine 6-phosphate | 2.14E-02 | 7.22E-03 | 2.6 | 4.1 |
| N-Acetylglucosamine 1-phosphate | 2.20E-02 | 3.64E-03 | 1.8 | 3.0 |

**b**

|  | **pval.adj.chimp** | **pval.adj.mac** | **FC.chimp** | **FC.mac** |
| --- | --- | --- | --- | --- |
| AMP | 1.65E-02 | 9.30E-03 | 6.5 | 10.0 |
| IMP | 9.93E-04 | 1.87E-02 | 14.8 | 18.3 |
| GMP | 1.61E-02 | 9.30E-03 | 3.7 | 4.9 |
| UDP-N-acetylglucosamine | 2.02E-02 | 1.87E-02 | 2.7 | 2.4 |
| NAD+ | 1.65E-02 | 9.30E-03 | 3.7 | 4.9 |

**Supplementary Tables’ description**

**Supplementary Table 1.** The concentration of metabolites detected by CE-MS in five tissues of four humans. chimpanzees. and macaques.

**Supplementary Table 2.** Metabolites showing higher concentration in human cerebellum **(a)** and lower concentration in human prefrontal cortex **(b)**. Table includes corrected p-values and fold-change for human-chimpanzee and human-macaque comparisons.

**Supplementary Table 3 a.b.** Pathway enrichment in metabolites showing **(a)** higher or **(b)** lower concentration in human tissues. Number of metabolites of **(a)** higher or **(b)** lower concentration and number of associated genes are indicated for each pathway. Top pathways (corrected p-value < 0.05) are colored in green.

**Supplementary Table 4.** Sample meta data table of the nine and eight tissues analyzed from wild type and humanized mice at one week and 12 week of age. Outlier samples are in red.

**Supplementary Table 5 a.b.** The concentration of metabolites detected by GC-MS in **(a)** eight tissues for 1 week- and **(b)** nine tissues for 12 weeks-old humanized and wild-type mice.

**Supplementary Table 6.** List of 45 metabolites showing significant differences in the prefrontal cortex (PFC) of one-week-old mice and list of 36 metabolites showing significant concentration differences in cerebellum (CB) of 12-weeks-old mice.
